# Whole-genome duplication reshapes adaptation: autotetraploid *Arabidopsis arenosa* leverages its high genetic variation to compensate for selection constraints

**DOI:** 10.1101/2025.04.01.646563

**Authors:** Sonia Celestini, Veronika Lipánová, Jakub Vlček, Filip Kolář

## Abstract

Whole-genome duplication (WGD), a widespread macromutation across eukaryotes, is predicted to affect the tempo and modes of evolutionary processes. By theory, the additional set(s) of chromosomes present in polyploid organisms may reduce the efficiency of selection while, simultaneously, increasing heterozygosity and buffering deleterious mutations. Despite the theoretical significance of WGD, empirical genomic evidence from natural polyploid populations is scarce and a direct comparisons of selection footprints between autopolyploids and closely related diploids remains completely unexplored. We therefore combined locally sampled soil data with resequenced genomes of 76 populations of diploid–autotetraploid *Arabidopsis arenosa* and tested whether the genomic signatures of adaptation to distinct siliceous and calcareous soils differ between the ploidies. Leveraging multiple independent transitions between these soil types in each ploidy, we identified a set of genes associated with ion transport and homeostasis that were repeatedly selected for across the species’ range. Notably, polyploid populations have consistently retained greater variation at candidate loci compared to diploids, reflecting lower fixation rates. In tetraploids, positive selection predominantly acts on such a large pool of standing genetic variation, rather than targeting *de novo* mutations. Finally, selection in tetraploids targets genes that are more central within the protein– protein interaction network, potentially impacting a greater number of downstream fitness-related traits. In conclusion, both ploidies thrive across a broad gradient of substrate conditions, but WGD fundamentally alters the ploidies adaptive strategies: tetraploids leverage their greater genetic variation and redundancy to compensate for the predicted constraints on the efficacy of positive selection.

## Introduction

Understanding how genetic variation and environmental pressures interact to shape species adaptation over time is a central challenge for research in our rapidly changing world (Rausher and Delph 2015). Among the many genomic processes influencing evolutionary dynamics, whole-genome duplication (WGD, i.e. polyploidization) emerges as a substantial and recurrent macromutation with far-reaching, yet still under-explored, implications for evolution and ecology (Fox et al. 2020). WGD is widespread in the Eukaryotic kingdom and has significantly affected the evolutionary history of plants in particular. Indeed, all plants are paleo-polyploids, meaning that they all have experienced at least one episode of polyploidization (Wendel 2015; Alix et al. 2017), and multiple ancient WGDs have been recognized in evolutionary history of land plants (Garsmeur et al. 2014; Zhang et al. 2020).

Earlier ideas saw polyploidy as an evolutionary ‘dead-end’ because of the expected long-term detrimental effects of increased genomic complexity on competition and adaptation (Comai 2005; Wendel 2015; Van de Peer et al. 2017). Nevertheless, several ancient polyploidization events seem to coincide with major lineage diversifications (Van de Peer et al. 2017), adaptive radiations (Zhang et al. 2020), enhanced invasiveness and colonization potential (te Beest et al. 2012; Baduel, Bray, et al. 2018; Baniaga et al. 2020), and plant domestication success (Paterson 2005; Salman-Minkov et al. 2016). The abundance and observed adaptability of polyploids in nature thus reinforces the more recent view of WGD as an advantageous opportunity for evolutionary success (Baduel, Bray, et al. 2018). However, the consequences of WGD on natural selection processes remain controversial (Soltis and Soltis 2000; Comai 2005; Parisod et al. 2010; Carretero-Paulet and Van de Peer 2020). To disentangle this conundrum, the study of autopolyploidy, where genome doubling happens within a single species, helps to isolate the effect of polyploidization from the confounding factor of hybridization present in allopolyploids. Empirical population genomic studies on autopolyploid species are, however, rare (Konečná et al. 2021; Bray et al. 2024; Hämälä et al. 2024; Zhang et al. 2024) and we completely lack a systematic analysis that would compare positive selection footprints in diploid and closely related autopolyploid genomes facing similar selective pressure.

Recent theoretical exploration on quantitative and population genetics of autopolyploidy has led to an appreciation of the potential dual role of WGD in shaping the tempo and modes of evolution in the adaptive landscape (Cuypers and Hogeweg 2014; Baduel, Bray, et al. 2018; Yao et al. 2019; Monnahan and Brandvain 2020; Clo 2022; Ebadi et al. 2023). On the one hand, the additional set of chromosomes in autopolyploid organisms masks recessive alleles in a heterozygous state and thereby, theoretically, increases the genetic load (Ronfort 1999) and diminishes the efficiency of purifying and positive selection, slowing fixation (Monnahan and Brandvain 2020). On the other, however, the increased heterozygosity in autopolyploids can buffer the effects of deleterious mutations and thus, potentially enhance genetic flexibility (Parisod et al. 2010). Here, constraints on gene evolution relax and, contrary to what is generally observed in plants (Masalia et al. 2017), fast adaptation might happen thanks to selection acting on proteins that are central in molecular networks, meaning that they affect a greater number of downstream fitness-related traits (Wang et al. 2010; Frachon et al. 2017; Rennison and Peichel 2022). Additionally, weaker genetic linkage and increased effective recombination rates (e.g. Monnahan et al. 2019), which result from the higher number of chromosome combinations at meiosis, theoretically reduce Hill–Robertson interference (Hill and Robertson 1966) and unlink unfavourable allele combinations more efficiently. In simulated polyploid genomes, higher recombination rates and extended fixation times due to masking create a unique genomic footprint, distinct from what is expected in diploids (Monnahan and Brandvain 2020). Hard selective sweeps in polyploids cause a greater absolute reduction in diversity (sweep magnitude) compared to diploids but remain more localized (narrower breadth) and do not reach fixation. However, empirical evidence from natural populations supporting these expectations is missing.

The dynamics, speed and outcome of adaptation are affected by the source of variation – standing or *de novo* – targeted by natural selection (Barrett and Schluter 2008; Thompson et al. 2019), and this has been demonstrated by ample genomic evidence in diploids (Reid et al. 2016; Alves et al. 2019; Ji et al. 2020; Bohutínská et al. 2021; Choi et al. 2021; Montejo-Kovacevich et al. 2022; Rubin et al. 2022). However, the role of WGD in modulating sources of adaptive variation in natural populations have never been addressed. Konečná et al. (2021) found repeated instances of adaptation to toxic substrates in autotetraploid populations of *Arabidopsis arenosa* originating predominantly from standing variation. However, the lack of comparable environmental contrasts in diploid populations left the role of ploidy unclear. At the same census population size, tetraploid populations are expected to experience close to double the occurrences of *de novo* mutations compared to diploids, due to a doubled number of mutation targets (Selmecki et al. 2015). Whether individual polyploid populations differently leverage such novel genetic variants or instead rely on the high level of standing genetic variation already present in the polyploid lineage can markedly affect their adaptive success. Taken together, all these genetic processes have influenced the past evolutionary trajectories of polyploids and will affect their evolvability and future survival in a fast-changing world (Wu et al. 2020). However, the balance between the advantages and disadvantages of WGD in shaping genetic adaptation remains a critical aspect that remains to be addressed empirically.

In this study, we address this knowledge gap by investigating how diploid and closely related autotetraploid populations of *Arabidopsis arenosa* adapt to non-extreme, spatially heterogeneous soil factors commonly experienced by plants, namely those determined by calcareous and siliceous substrates. Soils of these types impose different sets of challenges for plant life, especially in terms of pH and nutrient availability (Ormeño et al. 2008; Bontpart et al. 2024). They are both frequently inhabited by *A. arenosa* throughout central and southeastern Europe (e.g. Morgan et al. 2020 and Guggisberg et al. 2018) has detected related signatures of adaptation, in a pooled data of seven diploid populations, in genes involved in ion transport, response to nitrate and cellular ion homoeostasis. Here we developed on these successful findings and expanded the dataset to individually sequenced diploid and autotetraploid genomes of plants from 76 populations covering the range-wide diversity of *A. arenosa*. We complemented each population sample with *in situ* sampled soil data and applied a set of methods inferring positive selection to unravel the genetic basis of substrate adaptation in each ploidy level. Then, by comparing the top candidate selection sweeps across ploidies, we sought to answer the following questions: i) Do tetraploids rely on novel private (post-WGD) variation or rather adapt through the same genes selected for also in diploids when facing the same selective pressure? ii) Is there a larger proportion of fixed variants in diploids compared to tetraploids? iii) Do selection footprints differ between ploidies in terms of sweep breadth and magnitude? iv) Do ploidies differ in their ability to repeatedly leverage distinct sources of adaptive variation (standing variation, migration, *de novo* mutations)? v) Does positive selection in tetraploids target genes that are more evolutionary constrained (i.e. more central in the protein–protein interaction network) than are candidate genes inferred in diploids?

## Results

### Siliceous and calcareous soils were colonized repeatedly by each ploidy

We collected genomic data for 76 populations (35 diploids, 2x, and 41 tetraploids, 4x; 479 individuals in total), covering the majority of the distribution range of *Arabidopsis arenosa* and different soil types (Fig. 1a). Analyses of locally collected soil samples identified soil pH, calcium availability and cation exchange capacity (CEC) as the major differentiating variables (PC1 explains 44% of the variation, Fig. 1c). As expected, samples of calcareous soils (here defined by PC1 scores smaller than < 0) exhibit high pH, large concentrations of exchangeable calcium, generally high CEC and lower concentrations of potassium, which are typical characteristics of calcareous conditions. In turn, samples of siliceous soils (PC1 score > 0) exhibit the opposite trends. Note that other potentially relevant environmental factors, like temperature and precipitation, do not generally associate with soil conditions (supplementary fig. S1). In addition, neither soil environmental conditions, nor climatic parameters differ between the ploidies, as both ploidies span the broad range of environmental conditions (Fig. 1c, supplementary fig. S1), in line with previous reports of a lack of ploidy-related niche differentiation (Morgan et al. 2020; Padilla-García et al. 2023).

**Fig. 1.**
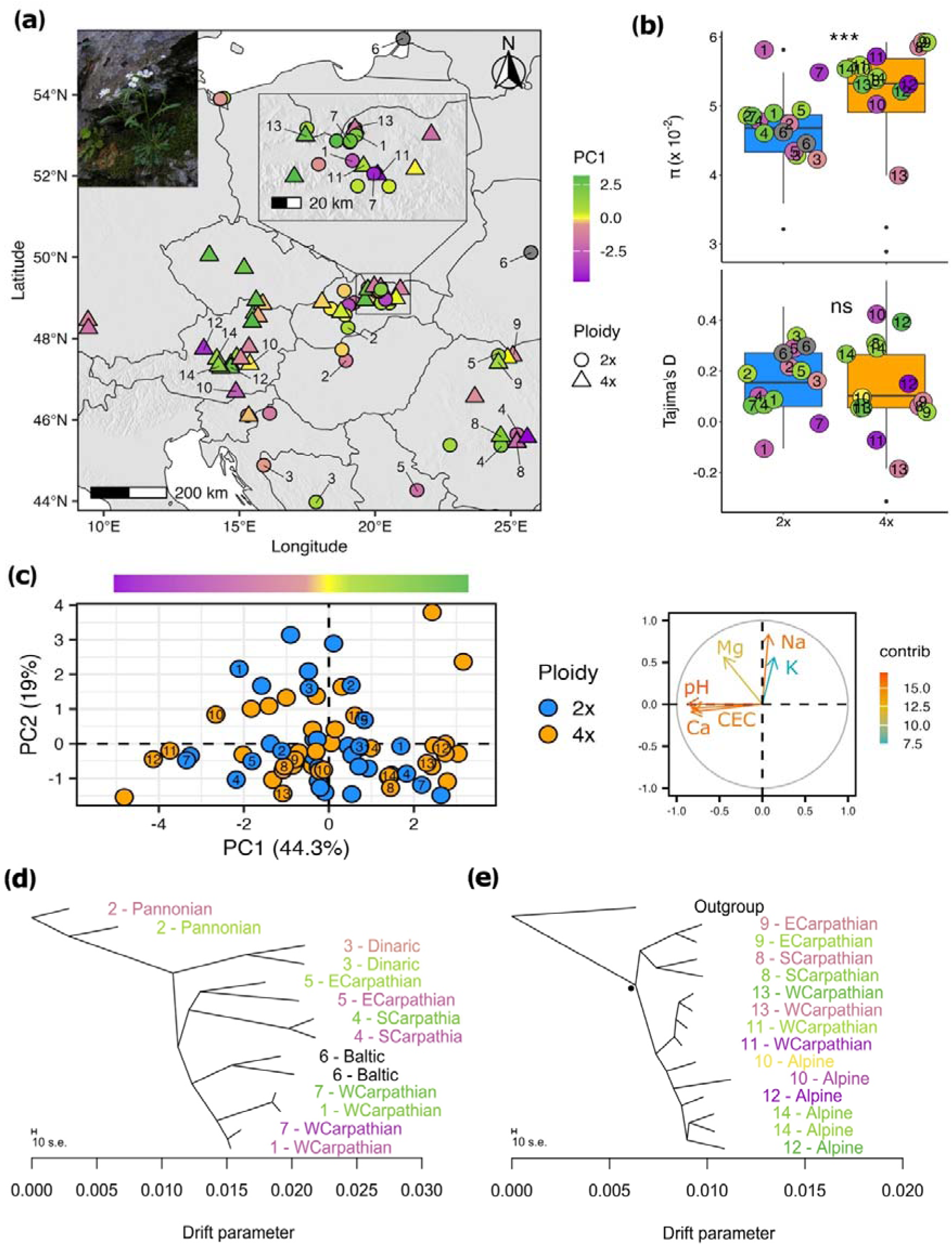
Genetic relationships, diversity, distribution and soil conditions of the *Arabidopsis arenosa* populations under study. a) Geographical distribution in central and southeastern Europe of the 76 genome-sequenced populations studied. Circles indicate diploid populations and triangles indicate tetraploid populations. The subset of populations used for pairwise divergence contrasts (paired dataset) are annotated with their relative pair number (diploid, 2x, 1–7; tetraploid, 4x, 8–14). b) Within-population diversity (pairwise nucleotide diversity, p) and estimate of neutrality (Tajima’s D) calculated from putatively neutral fourfold degenerated SNPs detected in 35 diploid and 41 tetraploid populations. Dots refer to the values for each population used in the paired dataset. c) PCA of the elemental soil composition for each population sampled (dots coloured by ploidy) (left) and variable correlation plot (right). Populations used in the paired dataset are indicated by their corresponding number. Soil variables are coloured by their contribution, expressed in percentage values, to the explained variation along the first two principal component axes. d,_e) Relationships between diploid (d) and tetraploid (e) populations of the paired dataset, depicted using an allele frequency covariance graph calculated in Treemix. Each population is labelled by its corresponding pair number and corresponding major genetic lineage, following previous range-wide studies (Monnahan et al. 2019; Padilla-García et al. 2023). All branches are significantly supported (bootstrap > 96%), the sole exception being marked with a black dot. The green–violet colour ramp reflects the position of each population along the PC1 axis of elemental soil composition, ranging from calcareous (violet) to siliceous (green) throughout the entire figure.

We investigated the genetic relationships between the populations and the possible repeated colonization of different soil types, using a set of 1,347,998 putatively neutral fourfold degenerate SNPs called in the full dataset of 479 *A. arenosa* individuals with an average sequencing depth of 25D. The principal component analysis (PCA – supplementary fig. S2) and the tree-based approaches (TreeMix – supplementary fig. S3, Fig. 1d, e), as well as the clustering method (Entropy – supplementary fig. S4), corresponded with the previously described internal sub-structuring of *A. arenosa* into distinct diploid and tetraploid lineages (Kolář et al. 2016; Monnahan et al. 2019; Padilla-García et al. 2023). The overall population grouping by spatial proximity and not by substrate indicates repeated colonization of contrasting soils by multiple lineages within each ploidy cytotype (Fig. 1d, e for the paired dataset, supplementary fig. S2–4 for the full dataset), albeit with an unclear ancestral state. Diploid and tetraploid populations show similar genome-wide Tajima’s D (average D_2x_ = 0.157, D_4x_ = 0.139; Welch two-sample t-test t(51.376) = 0.426, P = 0.672) that is close to neutrality, minimizing any possible effect of ploidy-specific demographic history on detecting signals of selection (Fig. 1b). In line with theoretical expectations (Baduel, Bray, et al. 2018), tetraploid populations show greater within-population nucleotide diversity than diploids (average π_2x_= 0.046, π_4x_ = 0.052; Wilcoxon rank sum test W = 166, P = 0.0005403) (Fig.1b).

### Genomic regions associated with adaptation to siliceous and calcareous soil differ between ploidies but share similar functions

We identified candidate genomic regions associated with genetic differentiation between populations from calcareous and siliceous soils for each ploidy level. To reduce the number of false positives, we used three complementary approaches and their intersection to refine and validate our list of top candidate *loci* for each cytotype. In addition, we focused on differentiation patterns recurrently present across the *A. arenosa* range and thus more likely to be result of selection as opposed to local (population pair-specific) signatures of genetic drift.

First, we scanned the genome for the top 1% of *F*_ST_ outliers among lineages growing on calcareous versus siliceous soil, calculated in 1-kbp windows including a minimum of 10 SNPs, in a subset of populations representing distinct genetic lineages of each ploidy (paired dataset, Fig. 1). Based on information on overall population structure and soil analyses, we selected seven ‘calcareous/siliceous’ population pairs (contrasts) per ploidy, maximizing the divergence in soil properties while minimizing neutral genome-wide differentiation and geographical distance within the pair (see the Methods section for details). This analysis resulted in an average of 564 and 643 outlier SNP windows in diploid and tetraploid pairs, respectively. After annotating the outlier windows to genic regions, we detected an average number of 406 annotated genes in diploids and 426 in tetraploids (supplementary data S2). We then identified a total of 233 genes in diploids and 219 genes in tetraploids that were outliers in at least two population pairs of different lineage within ploidy (Fig. 2c,_e). We refer to these genes as parallel candidate genes. Notably, these *loci* do not cluster in regions characterized by extreme recombination rates, indicating no bias associated with the recombination landscape (the average recombination rates in regions found to be under selection do not differ from the average recombination rate for genes not subject to selection, one-sided permutation test p-value = 0.9426). These genes are significantly enriched (p < 0.05) for an array of biological processes related to substrate adaptation such as xenobiotic transmembrane transport, cellular response to nitrate, potassium ion transmembrane transport, and lateral root development (supplementary data S3). Second, repeated genetic differentiation between populations growing on calcareous versus siliceous soil from the paired dataset was further quantified with PicMin (Booker et al. 2023), a statistical method which uses order statistics to quantify the non-randomness of adaptation across multiple lineages, providing a statistical confidence value for each SNP window. In this way, we identified 195 and 131 significantly differentiated 1-kbp windows (FDR-corrected q-value < 0.05) (Fig. 2a,_b), overlapping with 120 and 69 annotated genes in diploid and tetraploid populations, respectively (Fig. 2c,_e). Third, we leveraged the locally sampled elemental soil composition data and tested for any association between these data and SNP allele frequencies in the 33 diploid and 41 tetraploid populations for which we had gathered local soil chemistry data (see the Methods section for details) using latent factor mixed models (LFMM). Using population scores on the soil-associated PC1 axis to summarize the main gradient between siliceous and calcareous soils, we identified 817 (0.06%) and 515 (0.04%) SNPs that were significantly associated with soil PC1 (FDR-corrected q-value < 0.1) in diploids and tetraploids, respectively (Fig. 2a,_b). These SNPs overlapped with 33 and 22 annotated genes in diploids and tetraploids, respectively (Fig. 2c,_e).

**Fig. 2.**
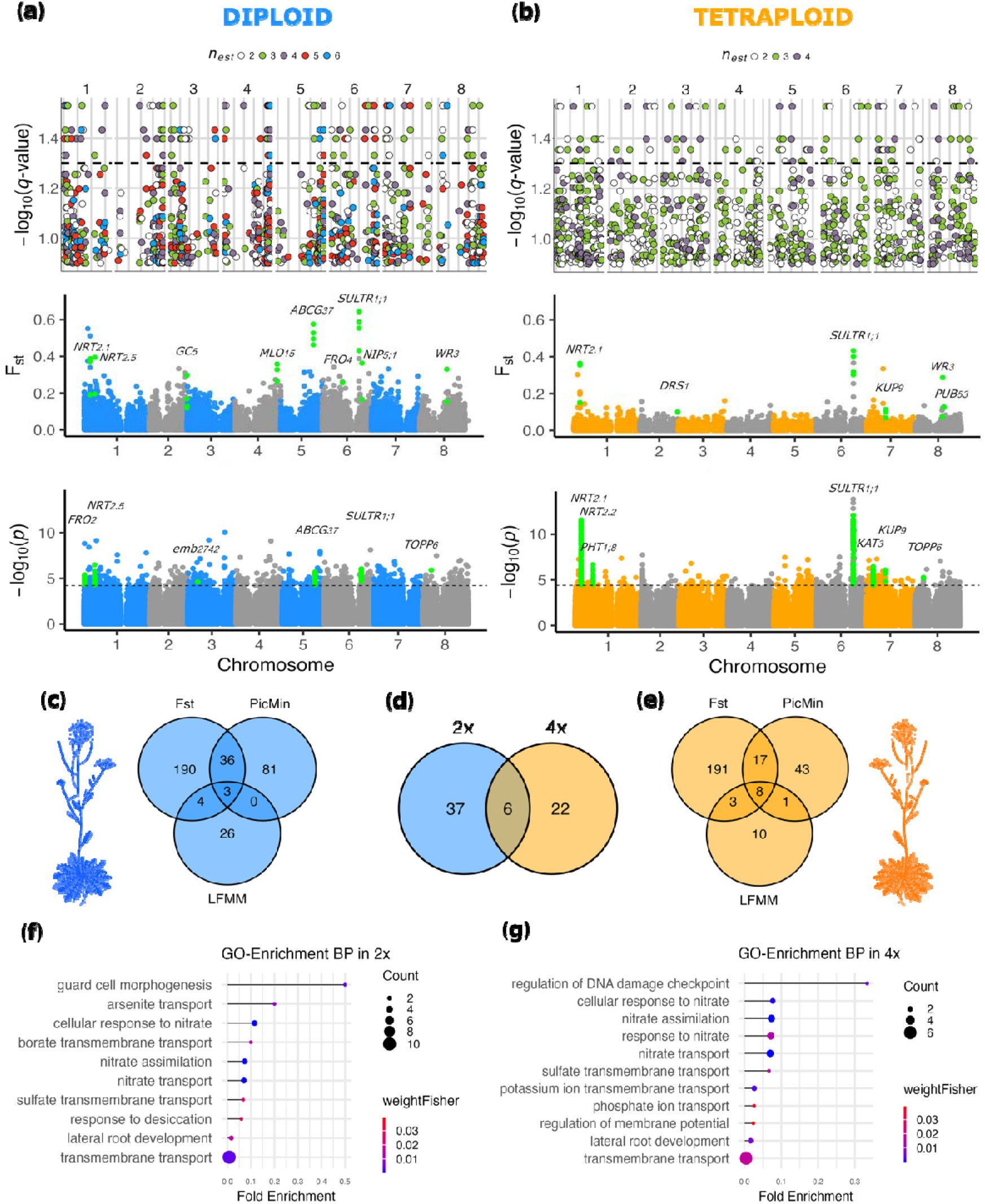
Genomic basis of calcareous/siliceous substrate adaptation in diploid and autotetraploid *Arabidopsis arenosa*. a,_b) Manhattan plots comparing the differentiation between populations on siliceous versus calcareous soils in terms of: repeated differentiation evaluated by means of order statistics in PicMin (upper row), Weir and Cockerham *F*_ST_ per SNP (middle row) and the association of per-SNP allele frequency and soil properties, approximated by scores on the soil-associated PC1 axis by means of latent factor mixed model (LFMM) analysis (lower row), in diploid (a) and tetraploid (b) populations. In PicMin, the dashed line indicates a significance threshold of *q* = 0.05, inferred by means of order statistics, and *n*_est_ refers to the estimated number of contrasts (i.e. the couples used in the analyses) exhibiting a pattern of repeated adaptation. In the LFMM analysis, the black dashed horizontal line refers to the applied threshold calculated as the 0.05 *q*-value. Selected top gene candidates with known function relevant to substrate adaptation are highlighted in green and annotated with their name. c,_e) Venn diagrams for diploid (c), and tetraploid (e) describing the total number of candidate genes overlapping between the three methods applied and their overlap across ploidy. d) Venn diagram depicting the overlap of selected top genes between the ploidies. f,_g) Significantly enriched biological processes (BP) gene ontology (GO) terms identified among the top candidate genes in diploids (f) and tetraploids (g).

Finally, we overlapped the results between methods to produce a refined list of ‘top candidate genes’ (Fig. 2d), retaining only the genes identified by the *F*_ST_ selection scan and at least one of the other methods (43 genes in 2x and 28 genes in 4x). This is a rather conservative approach aimed to minimize false positive signals that might stem from population-specific evolutionary history and genetic drift. Consequently, our approach is not aimed to cover the complete list of candidate genes, as it primarily focuses on hard sweep signals and neglects footprints of local site-specific selection taking place in certain population pairs. The list of candidate genes identified by each method for each population pair are also published as supplementary data S2 and S4. The list of top candidate genes encompasses biological functions relevant to substrate adaptation, mostly shared between the ploidies, related to ion (nitrate, potassium, phosphate, sulfate, borate, arsenite) transport, assimilation and response, and lateral root development (Fig. 2f,_g, supplementary data S5 and S6). Six top candidate genes were found among both diploids and tetraploids, which was more than random overlap of the genes under comparison (Fisher’s exact test; P = 5.09e-12) (Fig. 2d, Table 1).

**Table 1.** Description of the six top candidate genes shared between ploidies.

| *A. lyrata* ID | *A. thaliana* ID | Other name | TAIR functional description (Berardini et al. 2015). |
| --- | --- | --- | --- |
| AL1G18450 | AT1G08090 | *NRT2.1* | High-affinity nitrate transporter. Up-regulated by nitrate. Functions as a repressor of lateral root initiation independently of nitrate uptake. |
| AL1G18460 | AT1G08100 | *NRT2.2* | Encodes a high-affinity nitrate transporter. |
| AL6G42430 | AT4G08620 | *SULTR1;1* | Encodes a high-affinity sulfate transporter. Expressed in roots and guard cells. Up-regulated by sulfur deficiency. |
| AL8G13770 | AT5G45050 | *WRKY16* | Encodes a member of the WRKY Transcription Factor (Group II-e) family. |
| AL8G16980 | AT5G43380 | *TOPP6* | Encodes a type I serine/threonine protein phosphatase. |
| AL8G25530 | AT5G51270 | *PUB53* | Involved in protein ubiquitination. |

### Selective sweeps differ by cytotype

To investigate footprints of selection as a function of ploidy, we further compared various metrics of global and local population differentiation in the sets of top candidate genes found for each ploidy. Genetic differentiation (Rho, suitable for comparison across ploidies – Ronfort et al. 1998; Meirmans et al. 2018) between tetraploid candidate genes was relatively more elevated, when divided by genome-wide differentiation, compared to diploid candidates (Wilcoxon rank sum test W = 231, P = 1.306e-05, Fig. 3a; we control for the effect of positive selection in supplementary fig. S6). By contrast, the average number of fixed variants per gene is significantly greater in diploid population contrasts (average for 2x = 2.94 and for 4x = 1.46). Whereas 26 top genes in diploids show at least one fixed SNP (60.47% of the total), in tetraploids only 5 (17.86% of the total) top genes show fixation across all population pairs (Fig. 3b). In line with this observation, the distribution of average allele frequency of SNPs from these top candidates (Fig. 3d) is shifted more towards intermediate values in tetraploids, while allele frequencies in diploid candidates are closer to fixation. In Fig. 4a and 4b we show this same trend, zooming in on the gene SULTR1;1, function of which is strongly relevant for substrate adaptation and previously described in Guggisberg et al. (2018). Note that this result is not influenced by the greater number of chromosomes sampled in tetraploid populations, as sub-setting the dataset to 12 chromosomes per population resulted in the same exact trend (i.e. a greater fixation rate in diploids, supplementary fig. S7).

**Fig. 3.**
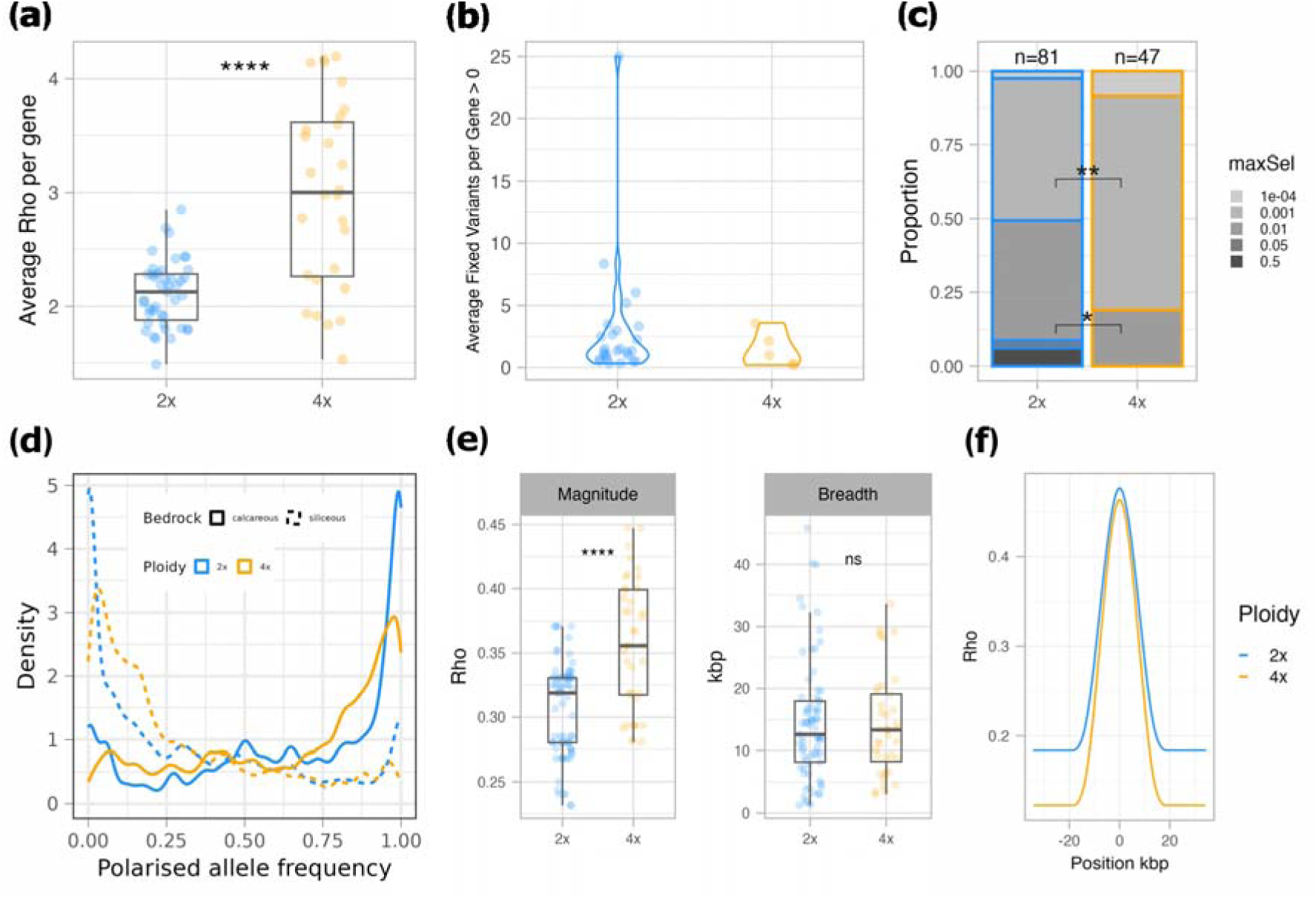
Different footprints of selection in top candidate genes for substrate adaptation in diploid and tetraploid *A. arenosa* populations (paired dataset). a) Average increase in genetic differentiation (Rho) in top candidate genes relative to genome-wide values. b) Average number of fixed variants per top candidate gene, calculated across overlapping 1-kbp sliding window and population pairs. In both a and b, each population pair was added to the calculation of the mean only if the gene was a significant outlier in that specific pair. c) Proportional representation of bins of selection strengths inferred by the DMC modelling method (see subsection) for each case of top gene parallel adaptation in each ploidy. Strength of selection (maxSel) is calculated as the most likely *s* at the identified selected gene site for the model with the highest composite log-likelihoods in DMC. We detected a significant association between ploidy and maxSel values of 0.001 and 0.01 using Fisher’s exact test (two-tailed P = 0.009533 and P = 0.01846, respectively). d) Distribution of average allele frequency per SNP (polarised by soil type to aid visualization in the functional context) across ploidies and soil types. Average allele frequency values were calculated per each SNP belonging to a top candidate gene, with a flanking region of 2 kbp. Only candidates from their respective population pairs where they appeared as outliers were included in the calculations. Note that the average allele frequencies in tetraploids are far from fixation and distributed more around intermediate values. e) Sweep magnitude and breadth for each top candidate gene across relevant population pairs. f) Smoothed-profile of the average selective sweep in each ploidy.

**Figure 4.**
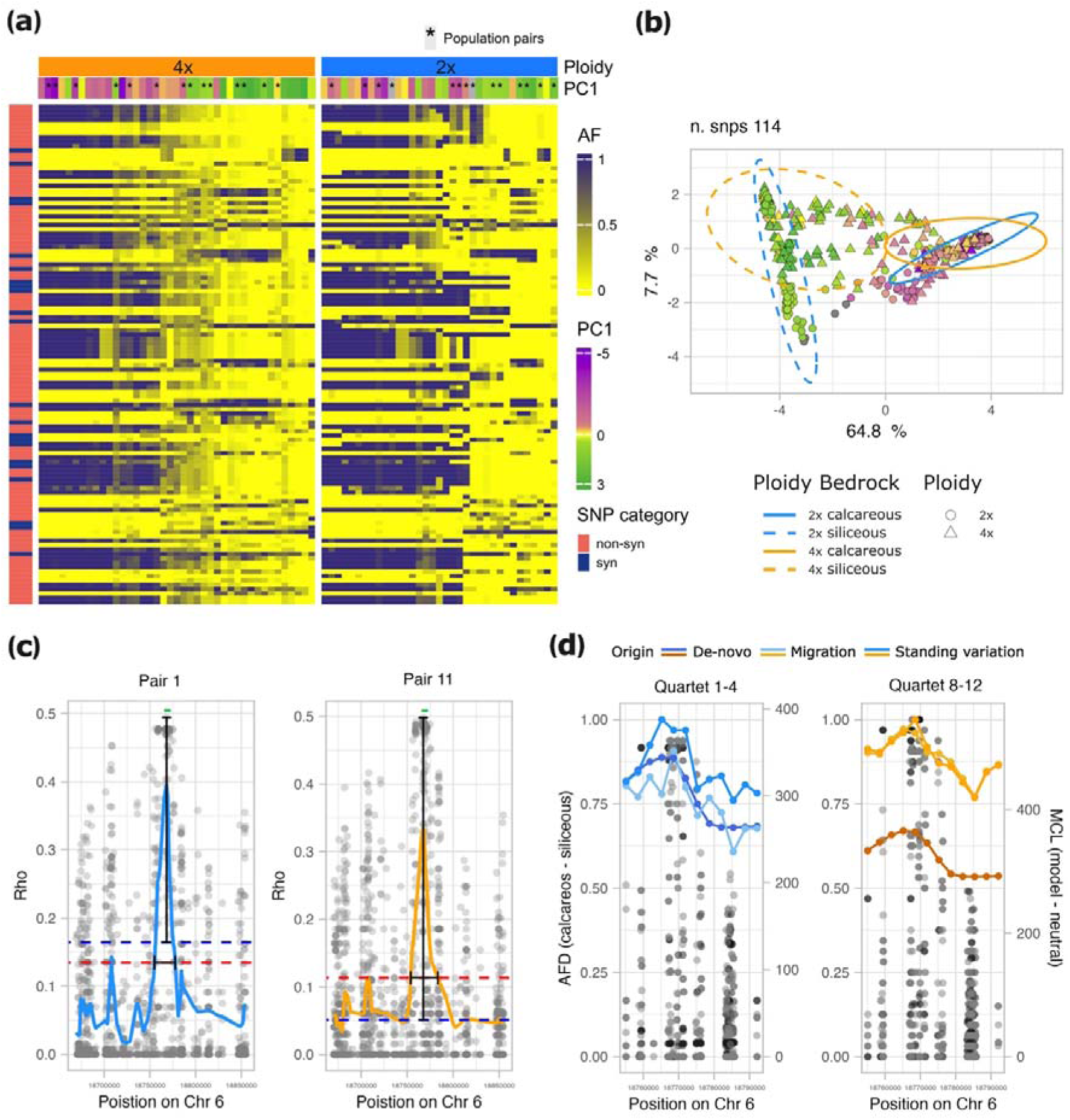
Different footprints of selection exemplified in one candidate gene found in both diploid and tetraploid *A. arenosa* (AL6G42430, SULTR1;1). a) Column-clustered heatmap showing allele frequency values for all 114 biallelic SNPs (rows) belonging to the gene (including UTRs, the transcribed region, exons and introns) per each population (column) of the full *A. arenosa* dataset. Colours in the top bars indicate the population ploidy and score along the soil-associated PC1 axis. Rows referring to synonymous and non-synonymous SNPs are coloured dark blue and red, respectively. In the heatmap, full purple cells indicate fixation of the ancestral allele whereas yellow cells indicate absence of the ancestral allele. Tetraploids exhibit a higher incidence of intermediate allele frequencies of individual sites values (0.5) than diploids. b) Local PCA calculated based on variation in the 114 SNPs of the SULTR1;1 gene. Each symbol refers to an individual and colours refer to the population score along the soil-associated PC1 axis. Note that whereas the calcareous/siliceous gradient explains most of the SNP variation (PC1 64.8%), geography, corresponding to scores on PC2, (not shown) has a markedly smaller effect. Tetraploid individuals are spread over a larger area in the plot and exhibit less differentiation between soil types, in line with the overall more intermediate allele frequency at the level of selected genes as compared to diploids. c) Selective sweep profile in diploid population pair 1 (left) and in tetraploid population pair 11 (right), indicated as a fitted curve along the distribution of differentiation (Rho) per each SNP (dots) in a genomic window of 200 kbp. The red dotted line represents the local background threshold (twice the average Rho in the 200 kbp flanking regions) and the blue dotted line represents the genome-wide average Rho. The green segment refers to the position of the SULTR1;1 gene model. The vertical black segment is the measured sweep magnitude, and the horizontal black segment is the measured sweep breadth. d) Allele frequency difference (AFD) between populations on calcareous versus siliceous soil for the locus (dots) and the maximum composite log-likelihood (MCL) estimation of the source of the selected alleles, inferred in DMC (lines), one of the diploid quartets on the left and one of the tetraploid quartets on the right. In both ploidies the standing variation origin scenario exhibits the highest likelihood.

Using the composite-likelihood-based model approach provided by the DMC pipeline (Lee and Coop 2017), we estimated the selection strength (*s*) acting on each top candidate gene (Fig. 3c). After filtering out cases which did not differ from a neutral scenario and retaining the results only for genes that were significant outliers in the respective set of populations, we obtained results for 81 cases (one candidate gene investigated in one quartet) in diploids and for 47 cases in tetraploids. In the latter, most of the cases (81%) concern genes affected by very low *s* (0.0001, 0.001) whereas in diploids, only 50% of the case show that same intensity of selection. Besides, genes showing high strength of selection (0.05, 0.5) are present only in diploids, where they represent 8% of cases. Further, we investigated if hard selective sweep features at the locations of candidate genes differed in each ploidy, following the expectations formulated in the simulation study of Monnahan and Brandvain (2020). For each case of selection (i.e. the combination of a top candidate gene and population pair(s) in which the gene has been identified as an outlier) we determined the values of sweep magnitude and breadth from SNP differentiation (Rho) values (see Fig. 4c for an example calculation for the SULTR1;1 gene). On average, sweep magnitude (i.e. the increase in differentiation at the selective peak above the genome-wide differentiation) is significantly greater in tetraploids (average for 4x = 0.357, average for 2x = 0.306, Welch two-sample t-test t(61.703) = −5.6475, P = 4.402e-07), while sweep breadth is comparable between the ploidies (average 4x = 14.217, average 2x = 14.493, Wilcoxon rank sum test W = 1455, P = 0.9784) (Fig. 3e,_f). These same general trends, represented in Fig. 3, are maintained when focusing only on the six top candidate genes shared between the ploidies (supplementary fig. S8).

### Tetraploid populations do not leverage *de novo* mutations for adaptation

Increased variation harboured in the tetraploid population and efficient masking of rare novel (*de novo*) recessive variants may in sum imply a greater importance of sampling from pre-existing, shared (standing or introgressed) variation during the adaptation process in tetraploid populations relative to diploids. To test this hypothesis, we quantified the importance of different evolutionary sources of adaptive variation using the ‘distinguishing among modes of convergence’ (DMC) approach (Lee and Coop 2017). The DMC method uses patterns of covariance in allele frequencies near the site subject to selection to identify the most likely evolutionary mode of repeated adaptation between two pairs of populations repeatedly adapting to the same environmental factor. First specifying a null model that accounts for population structure, we tested for three alternative models: (i) independent *de novo* mutations at the same locus, (ii) selection acting on shared standing variation, and (iii) sharing of beneficial alleles via gene flow between populations. For each ploidy, we ran the DMC analysis over all top candidate regions along each of 21 population quartets (pairs of population contrasts) using the same population contrasts as in the PicMin analysis (see the Methods section) to retain only population contrasts that belong to significantly distinct intraspecific lineages (Fig. 1e,_f; supplementary fig. S3). Selective signatures in 40 genes out of 43 in diploids and 22 genes out of 28 in tetraploids were significantly different from neutrality and were therefore further considered in the analysis. In both ploidies, adaptation was shown to mostly originate from standing variation, where 54.32% (n = 44) and 59.57% (n = 28) of cases (genes per quartet) in diploids and tetraploids, respectively, were attributed with the greatest likelihood (based on maximum composite log-likelihood, MCLs) to the standing variation scenario, and this difference was not statistically significant between the ploidies (*X*^2^(1, N = 128) = 0.15, P = 0.6945) (Fig. 5a, supplementary data S7). However, whereas in diploids the percentage of parallel *de novo* origin scenarios was relatively high (18.52%, n = 15), in tetraploids this scenario was supported by the highest MCL in only one case (2.13%, n = 1, significant difference between the ploidies *X*^2^(1, N = 128) = 5.88, P = 0.015): gene SPINDLY (SPY) – AL3G23090 in quartet 8–9. In tetraploid populations, the increased sharing of adaptive variants through migration (38.3%, n = 18 in 4x; 27.16%, n = 22 in 2x) seems to compensate the lack of selection acting on *de novo* mutations, although the proportion of loci shared through migration is not significantly greater than in diploids (*X*^2^(1, N = 128) = 1.24, p = 0.2659).

**Figure 5.**
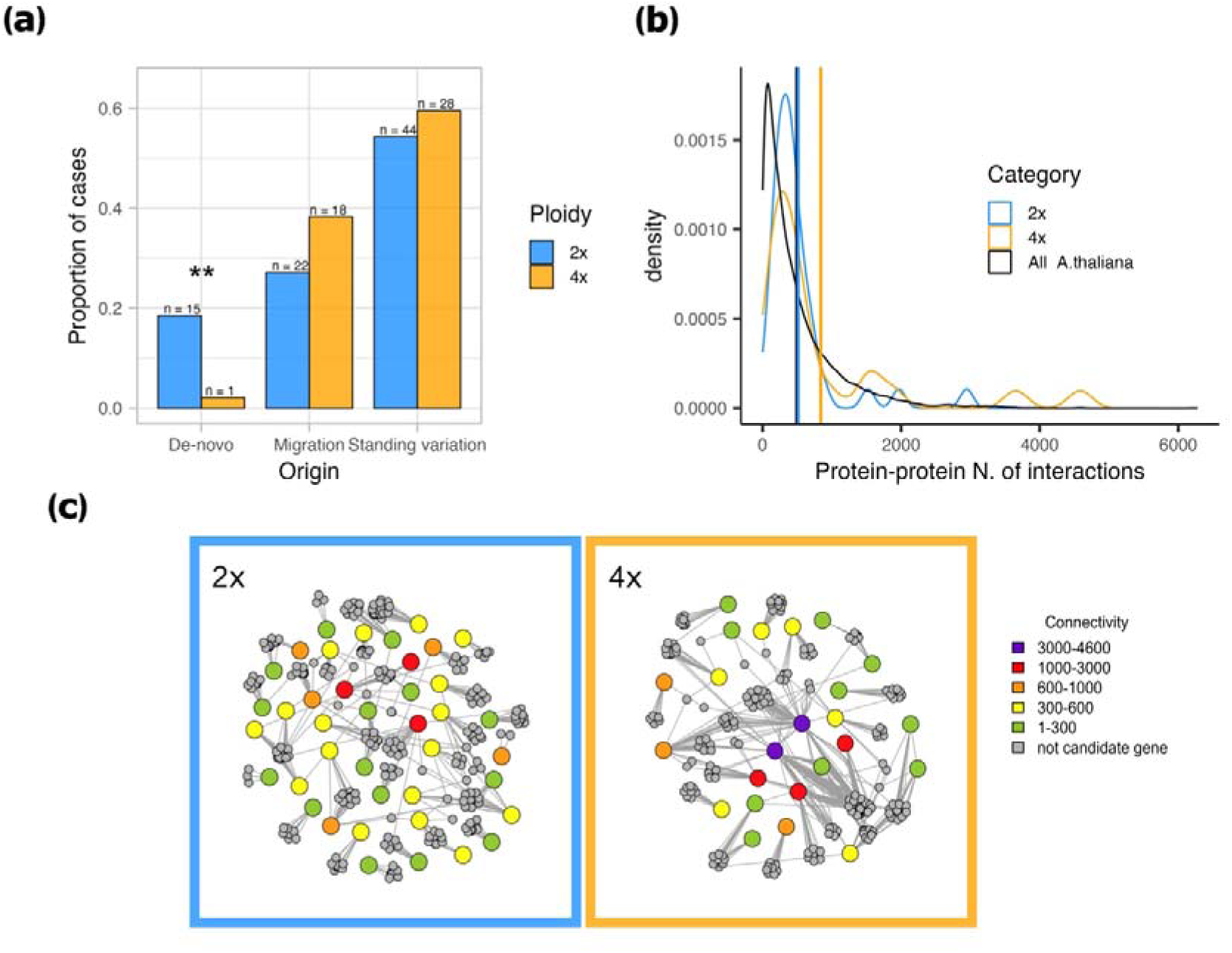
Different evolutionary sources and number of protein–protein interactions of candidate genes associated with substrate adaptation in diploid and tetraploid *A. arenosa* populations. a) Different representation of the three major evolutionary sources of repeated adaptation quantified by DMC in each ploidy. b) Density distribution of the number of protein–protein interactions among the top candidate genes per each ploidy, compared to the full dataset of *A. thaliana* genes. The vertical lines show the position of the mean of each group. c) Protein–protein interaction networks of the top candidate genes demonstrating overall greater connectivity (red/violet shading) of genes identified in tetraploids (right).

### Tetraploid adaptive candidates are more central in the protein–protein interaction network

Efficient and faster adaptation might be achieved in tetraploids if selection acts preferentially on genes that exhibit a central position in the protein–protein interaction network and thus likely have a strong pleiotropic effect (Promislow 2004). We measured this using the STRING database v.12 (https://string-db.org) (Szklarczyk et al. 2023) of the protein–protein interaction network in *Arabidopsis thaliana*. In total, we collected data for 25,715 *A. thaliana* proteins (nodes) annotated to 26,206 genes. The number of edges per connected node ranged from 1 to 6,278, with a mean value of 487.9 edges per node (Fig. 5b). We compared these values with the nodes’ connectivity measures for *A. thaliana* orthologs of our set of 40/24 top candidate genes in diploid/tetraploid *A. arenosa*. In diploids, the number of edges per node ranged from 34 to 2,947, with a mean value of 514.6; in tetraploids the range spanned from 107 to 4,587 edges per node, with a mean of 841 (Fig. 5b, supplementary data S5). The average connectivity of diploid top candidate genes does not significantly differ from the average number of edges in the complete *A. thaliana* dataset (one-sided permutation test P = 0.3989), indicating that candidate adaptive genes are neither more peripheral nor central than the average. By contrast, tetraploid top candidate genes show a significantly greater number of edges per node, and thus connectivity, than the average for *A. thaliana* (one-sided permutation test P = 0.0049).

## Discussion

### Repeated colonization of siliceous and calcareous substrates across the range of *A. arenosa* recruits genes involved in ion transport and homeostasis

Siliceous and calcareous soils are usually scattered throughout the landscape, creating a mosaic of contrasting habitats differing in pH, chemistry and texture. These characteristics are known to affect plant distribution and productivity worldwide (Michalet et al. 2002; Alvarez et al. 2009; Bothe 2015; Nemer et al. 2021; Smyčka et al. 2022). In many plant groups, differentiation into populations growing on calcareous or siliceous soils coincided with speciation (e.g*. Minuartia*, *Androsace*, *Campanula*, *Gentiana*, *Phyteuma*, *Primula* and *Saxifraga*) (Moore and Kadereit 2013; Smyčka et al. 2022; Lipánová et al. 2023). However, lineages and cytotypes of *A. arenosa* have repeatedly colonized both types of environments without accumulating significant genome-wide differentiation (Fig. 1). Such pattern rather suggests a scenario of repeated ecotypic differentiation similar to that observed in other plant species with variable substrate preferences (Bastida et al. 2014; Kolář et al. 2014) or in *A. arenosa*, populations adapted to high elevations (Bohutínská et al. 2021) and toxic serpentine soils (Arnold et al. 2016; Konečná et al. 2021).

Living on siliceous and calcareous soil requires specific mechanisms for dealing with many chemical constraints, such as high levels of calcium and carbonate, deficiencies of iron, zinc and potassium in calcareous soils, and with aluminium toxicity, nitrogen deficiency and decreased basic cation concentrations in siliceous soils (Clark and Baligar 2000), which may act as selective factors. Indeed, we observed enrichment for genes involved in ion transport and homeostasis (see below for a detailed discussion) among outliers identified in our analysis. This result complements previous findings made in a subset of diploid populations (Guggisberg et al. 2018), altogether suggesting an adaptive genetic response of *A. arenosa* to contrasting soil conditions. In line with these expectations, local adaptation to calcareous soil has been experimentally documented in the closely related species *A. lyrata* (Guggisberg et al. 2018) and *A. thaliana* (Terés et al. 2019), indicating that calcareous adaptation is a frequent phenomenon in the model genus *Arabidopsis* (Koch 2018).

As also described in Guggisberg et al. (2018), we identified the gene SULTR1;1 as a very strong adaptive candidate in both ploidies (Fig. 2 and 4). This gene is expressed in root epidermis and root hairs in response to sulfur starvation, and its up-regulation positively correlates with the content of sulfate in the roots of *Arabidopsis thaliana* (Takahashi et al. 2000; Rouached et al. 2009). Notably, sulfur deficiency is common in calcareous soils, as sulfur reacts with the abundant calcium present in the substrate, forming calcium sulfate (CaSO_4_), which prevents sulfur uptake. Yet, sulfur is a pivotal macronutrient for plants, affecting not only their growth and development, but also their response to oxidative stress, pests, diseases and heavy metals. Indeed, sulfur is required for the biosynthesis of compounds such as the amino acids cysteine (Cys) and methionine (Met), proteins, glutathione (GSH) and phytochelatins, vitamins, and secondary metabolites such as glucosinolates (Shah et al. 2022). One soil-specific SULTR1;1 allele has been found to be shared among 72 *Arabidopsis* accessions, suggesting that selection has recurrently acted on an ancient polymorphism existing in plants growing on soils of each type (Guggisberg et al. 2018). This is in accordance with our finding that selection concerning the SULTR1;1 gene acts mainly on standing variation across lineages and ploidies of *A. arenosa* (DMC results, Fig. 4d, supplementary data S7). Similarly to sulfur, potassium has an antagonistic relationship with calcium, which inhibits its uptake by plants (York et al. 1953). Accordingly, we found candidate genes related to potassium deficiency in *A. thaliana.* It is the GC5 gene, which encodes a protein that functions as a structural component of the Golgi apparatus in diploids (Kang et al. 2004), while in tetraploids selection acted on KAT3 gene, encoding potassium channel subunits, and on the KUP9 potassium ion transmembrane transporter (Hedrich 2012; Šustr et al. 2024). Interestingly, in diploids we have also found candidate genes involved in iron uptake, such as FRO2, FRO4 and ABCG37, which are well characterized in *Arabidopsis* (Vert et al. 2003; Fourcroy et al. 2016; Anjali et al. 2021). Iron is indeed inaccessible to plants in calcareous conditions, as ferric oxides become more insoluble with every unit increase in pH (Kim and Guerinot 2007).

In addition, we detected a strong signal of selection acting on three out of seven genes of the NRT2 family of nitrate transporters in *Arabidopsis* (NRT2.1, NRT2.2 and NRT2.5). These three genes are mostly expressed in the roots and, when induced by nitrogen starvation, mediate high-affinity root NO_3_^−^ influx (Xu et al. 2024). Although nitrogen soil content does not seem to be particularly linked to any single soil type, microbial activity, responsible for ammonification, nitrification, denitrification and nitrogen fixation, is usually lower at lower pH, negatively affecting how plants absorb nitrogen (Bothe et al. 2006). Many of the above listed genes have been identified as candidates for adaptation to serpentine soil in *A. arenosa* and *A. lyrata*, siliceous and calcareous conditions in *A. lyrata* and *A. thaliana,* metalliferous soil in *A. halleri* and salt stress in *E. salsugineum* and *A. pumila*, suggesting that they might represent genetic hotspots for adaptation to soil conditions (Turner et al. 2010; Arnold et al. 2016; Guggisberg et al. 2018; Sailer et al. 2018; Yang et al. 2018; Preite et al. 2019; Terés et al. 2019; Konečná et al. 2021; Tran et al. 2021).

Finally, one important distinguishing feature of siliceous and calcareous soils is their water and heat holding capacity. Typical calcareous soils are fissured, allowing rainfall to rapidly drain away, exposing plants to water deficiency and higher temperatures. By contrast, siliceous soils have almost no fissures and therefore retain moisture for longer, also maintaining a lower soil temperature. In agreement with this observation, four of our detected top candidate genes are enriched for drought resistance (RCI3, NIP5;1, DRS1, EXL4) (Llorente et al. 2002; Lee et al. 2010; Yu et al. 2015; Mancini et al. 2022). In summary, we have detected repeated signals of selection acting on a suite of genes affecting ion transport and management as well as drought resistance, all of which correspond with known challenges posed by the calcareous/siliceous soil gradient.

### Targets and strength of positive selection differ between ploidies of *A. arenosa*

Theory and simulations have raised clear expectations regarding how autopolyploids differently respond to positive selection due to their lower variance in fitness, different manifestation of dominance and propensity to retain genetic variation (Monnahan and Brandvain 2020). We addressed these expectations by comparing candidate genes allele frequency distributions and the extent of gene sharing between diploid and autotetraploid *A. arenosa* populations repeatedly adapting to similar environmental pressures.

First, we found that only six top candidate genes were shared between the cytotypes (14% of diploid top candidates and 21% of tetraploid top candidates, Fig. 2d). The number is greater than what would be expected randomly; however, it is surprisingly low considering the broadly shared pattern of polymorphism in *Arabidopsis* (Novikova et al. 2018) and the rampant gene-flow between the ploidies (supplementary fig. S2 and S4, Monnahan et al 2019). There are several possible reasons why the two ploidies rely mostly on different adaptive genes in spite of a globally shared pool of variation in the species. The first reason is that selection may act upon a different spectrum of allele dominance in each ploidy cytotype (Monnahan and Brandvain 2020). Fixation of recessive variants is expected to be less likely in polyploids due to much rarer frequencies of homozygotes (equilibrium *q*^4^) compared to diploids (*q*^2^). Selection in tetraploids might therefore be more efficient in genes where the adaptive variant is dominant, while diploids may rely on both types of variants (Monnahan and Brandvain 2020). Secondly, genes with different levels of centrality or genetic context could be differently accessible in higher or lower ploidies, which we discuss in more depth in the following section.

In addition, we have found a significant difference between the ploidies in the number of fixed candidate variants subject to selection and the characteristics of their hard selective sweeps. First, we detected an almost complete lack of fixed alleles in tetraploid populations (Fig. 3b), while selective sweeps are greater in magnitude than those of diploids; this means that the genetic differentiation between populations on siliceous and calcareous bedrock at the selected region is elevated higher above the genome-wide background in tetraploids (Fig. 3e,_f). In line with the reasoning explained above, we speculate that this might be a result of positive selection acting on mainly dominant variants in tetraploids. Indeed, simulations suggest that generally (partially) dominant mutations increase in frequency early on but then stabilize at intermediate frequencies and take longer to approach fixation; this is especially true in polyploid populations, where the frequency of the genotype expressing the dominant phenotype is higher than in diploids (Monnahan and Brandvain 2020). Moreover, in higher ploidies, time to fixation is expected to be longer also because the level of diversity before the onset of selection is usually higher (Otto and Whitton 2003). In turn, our results show that candidate alleles in diploids were more often fixed, which corresponds with the expected greater likelihood of selection acting on recessive alleles in lower ploidies (Marad et al. 2018; Monnahan and Brandvain 2020). An additional explanation lies in population-scaled recombination rates, which are expected to be higher in autotetraploids due to the additional chromosomal partners available during recombination (Pecinka et al. 2011). A higher population recombination rate could slow down the fixation of variants, promoting ‘haplotypic homogenization’ (high representation of intermediate allele frequencies at candidate loci). Yant et al. (2013) previously detected a lower-than-double crossover rate in autotetraploids of *A. arenosa*. This may explain why, in our analysis, sweeps breadth are similar between ploidies (Fig. 3d,_e). However, the greater sweep magnitude at similar sweep breadth in polyploids suggests a higher number of recombination events, potentially influencing fixation rates (Fig. 3d,_e). Finally, the observed ratio magnitude/breadth could also indicate stronger selection (*s*) in tetraploids. Yet, when *s* is estimated as the amount of decaying within-population haplotypic similarity around selected loci, tetraploid populations will exhibit lower apparent selection intensities during adaptation (as observed in our DMC results, Fig. 3c), which is consistent with an over-representation of intermediate allele frequencies in candidate loci.

### Sampling pleiotropic genes from shared variation facilitates fast adaptation, especially in tetraploid populations

Distinguishing among sources of adaptive variation (*de novo* mutations, gene flow or standing variation) provides insight about the evolutionary potential of populations, lineages or cytotypes. Theory suggests that access to sources of adaptation may differ between diploid and autopolyploid populations, but the direction of this difference remains equivocal (Van de Peer et al. 2017; Konečná et al. 2021). In the present study, we addressed this controversy by empirically inferring evolutionary sources of adaptive variation from naturally replicated instances of substrate adaptation using the modelling approach distinguishing among modes of convergent adaptation (DMC). We found that the common standing pool of variation ancestrally retained across populations of the same cytotype is the main source of adaptation in both ploidies (Fig. 5a). Predominant parallel adaptive sourcing from standing variation is in line with expectations for closely related populations (Bohutínská et al. 2021; Bohutínská and Peichel 2024) and has been documented in many diploid systems adapting to diverse challenges, such as vertebrates (Jones et al. 2012; Reid et al. 2016; Terekhanova et al. 2019; Louis et al. 2021; Rubin et al. 2022; Gutiérrez-Guerrero et al. 2024), insects (Ji et al. 2020; Chaturvedi et al. 2022; Montejo-Kovacevich et al. 2022) and plants (Bohutínská et al. 2021; Choi et al. 2021), but also autotetraploids of *A. arenosa* adapting to serpentine conditions (Konečná et al. 2021). However, we have found a significant difference in the relative importance of sources of adaptive variation between ploidies. In particular we found a virtual lack of tetraploid adaptation building on *de novo* mutations. This might reflect a strong effect of masking on novel mutations in tetraploids, as they are initially present in lower frequencies. In fact, a standing beneficial allele is more likely to be fixed, as it is likely to be present in a greater frequency at the time when selective conditions change (Innan and Kim 2004; Barrett and Schluter 2008). Moreover, standing variation is likely to lead to more rapid evolution in novel environments because beneficial alleles are immediately available, in contrast to new beneficial mutations, which require time to arise (Ji et al. 2020; Rubin et al. 2022). All else being equal, the fixation time of an allele is affected by the magnitude of its beneficial effect, the effective population size (*N*e) and the number of copies of the allele, which is inversely proportional to the likelihood of the allele being lost because of genetic drift (Hermisson and Pennings 2005). Consequently, considering that polyploid populations often display a higher *N*e and/or harbour a larger pool of variation (Van de Peer et al. 2017; Baduel, Bray, et al. 2018), polyploids should predominantly source their adaptation from pre-existing variation, including through possible between-population gene flow.

In addition to population genetic parameters, adaptation speed and success can be linked to the propensity of target genes, or their products, to undergo evolutionary changes, and it has been suggested that such a constraint depends on the gene’s position in the molecular network (Orr 2000). However, it remains unclear whether such characteristics differ depending on ploidy. In a protein– protein interaction network (PPIN), proteins occupying a central position have been shown to exert a strong influence over the pathway output, and thus on the final phenotypes and the organism’s fitness (therefore, connectivity correlates positively with pleiotropy – Promislow 2004; Tew et al. 2007; Tyler et al. 2009; Wang et al. 2010). Therefore, positive selection on genes encoding central proteins can drive rapid adaptation; however, here new mutations are more likely to be deleterious and are consequently quickly purged (Olson-Manning et al. 2012). Interestingly, in accord with theory (Wang et al. 2010) and recent empirical studies on sticklebacks (Rennison and Peichel 2022) and *Arabidopsis thaliana* (Frachon et al. 2017), we found that repeatedly selected candidate genes of *A. arenosa* have, on average, an intermediate level of pleiotropy/connectivity (Fig. 5b). Thus, this result adds to the existing body of work that conflicts with earlier theoretical expectations suggesting that higher levels of pleiotropy are constraining and disfavour adaptation (Otto 2004; Kim et al. 2007; Masalia et al. 2017). Notably, large-effect loci have been suggested to possess a selective advantage when a population is colonizing a new environment (Orr 1998; Hämälä et al. 2020), because the cost of complexity imposed by pleiotropy gets balanced out by its positive effect pushing the population faster towards a new fitness optimum (Wang et al. 2010). This process might even be less costly when selection targets adaptive loci standing in the population pool that are likely to have been pre-tested by selection during previous selective pressures (Barrett and Schluter 2008). Here, we suggest that the species *A. arenosa* as a whole might have been able to quickly adapt to new stress factors associated with siliceous or calcareous bedrock by leveraging intermediate-effect loci that were likely to be advantageous because sourced mainly from pre-tested standing or shared variation.

Moreover, candidate genes of the tetraploid cytotype display a significantly higher average level of connectivity than diploid ones (Fig. 5b,_c). Theory proposes that expression networks in polyploids are characterized by a greater redundancy given by the multiplication of each node (i.e. gene) and edge (i.e. interaction) (Parisod et al. 2010; Ebadi et al. 2023). Our results suggest that this redundancy, in line with simulations results from (Yao et al. 2019) might promote robustness against new mutations, allowing even more central genes to explore novel protein interactions and outputs while preserving their ancestral functions (Cuypers and Hogeweg 2014; Carretero-Paulet and Van de Peer 2020; Parisod 2024). For the reasons above, WGD has the potential to break genomic constraints imposed by pleiotropy, a mechanism that might compensate for the slower rate of fixation and the dominance-dependent efficiency of selection being experienced, allowing polyploids to ‘jump’ in the fitness landscape quickly reaching new fitness optima – hence the definition of ‘hopeful monsters’ (Cuypers and Hogeweg 2014; Keane et al. 2014; Ebadi et al. 2023). However, further empirical work directly comparing diploid *A. arenosa* against tetraploid gene regulatory networks would be beneficial in order to corroborate and shed light on the mechanisms of such a process.

## Conclusions

Here, we have identified and compared signatures of positive selection across ploidies, providing insights into how whole-genome duplication influences evolutionary trajectories. Our results support a scenario where, despite the potential constraints imposed by polysomic masking and the delayed fixation of beneficial alleles, tetraploid populations efficiently navigate selective pressures. Their inherent retention of a substantial reservoir of standing genetic variation translates into an evolutionary potential that enables them to persist and diversify in dynamic landscapes, be it in response to gradual ecological shifts or rapid environmental perturbations (Barrett and Schluter 2008; Keane et al. 2014; Fox et al. 2020).

## Materials and Methods

### Model species

*Arabidopsis arenosa* is an outcrossing species exhibiting very high genetic diversity enabling the efficient detection of hard selective sweeps (Yant and Bomblies 2017). Moreover, it has a well-characterized ploidy distribution, ecology and evolutionary history. Known autotetraploid populations of *A. arenosa* trace back to a single origin from the Western Carpathian diploid lineage ∼_19–31 k generations ago, after which they have spread across Europe, largely overlapping with the ecological niche of diploids, as the two ploidies followed a similar process of postglacial expansion (Arnold et al. 2015; Monnahan et al. 2019; Padilla-García et al. 2023). Notably, polyploid genomes have retained extensive genetic variation with no detectable effect of any conspicuous bottleneck, possibly reflecting a gradual origin from a large and diverse diploid population (Arnold et al. 2015). These factors make the age and demographic histories of populations involved in the colonization of different soils comparable between the ploidies and thus suitable for the purposes of our study. A pervasive effect of ploidy on genetic diversity and footprints of selection has been documented throughout the genome of *A. arenosa* (Monnahan et al. 2019). However, the observed trends primarily reflect genome-wide effects of neutral processes and purifying and linked selection, but the effect of ploidy on local signals of positive selection remains unknown.

### Genetic and soil dataset collection

Siliceous and calcareous soils exhibit a patchy distribution throughout the European landscape and ploidies of *Arabidopsis arenosa* occur in both these environments throughout their native ranges. Aiming to cover the majority of the species’ distribution along the soil gradients, we assembled whole-genome resequencing data for 76 populations (479 individuals) covering all lineages of *A. arenosa* (in line with Monnahan et al. (2019) and Padilla-García et al. (2023) except the invasively spreading Ruderal lineage, which exhibits weak substrate preferences, as it occupies human-disturbed non-native habitats (Baduel, Hunter, et al. 2018; Monnahan et al. 2019). Specifically, we used previously published data on 35 verified diploid (2x) and 41 verified tetraploid (4x) populations (Hollister et al. 2012; Yant et al. 2013; Arnold et al. 2016; Novikova et al. 2016; Monnahan et al. 2019; Bohutínská et al. 2021; Konečná et al. 2021), accessible under BioProjects PRJNA284572, PRJNA309929, PRJNA357693, PRJNA357372, PRJNA459481, PRJNA493227, PRJEB34247 (ENA), PRJNA506705, PRJNA484107, PRJNA592307, PRJNA667586, PRJNA929698 (see supplementary data S1 for details). At each population location, ∼_1 litre of soil was dug from ∼_10–20 cm below the ground in three different spots in the vicinity of *A. arenosa* plants. Two diploid Baltic populations included in this study lacked soil information because of complicated logistics (occurrence in Ukraine and Lithuania), but they were included in the selection scan (see below), reflecting their sharply contrasting substrate preferences inferred from our field notes on the type of bedrock and accompanying vegetation (a limestone outcrop vs acidic sandy soil).

### Soil analysis

Soil samples from each population were mixed, air-dried, sieved and analysed at the Analytical Laboratory of the Institute of Botany of the Czech Academy of Sciences in PrulJhonice following the protocol published in Konečná et al. (2021). The parameters measured were pH H_2_O, pH KCl, bioavailable Ca, total content of nutrients, such as Ca, Mg, K and Na, and cation exchange capacity (CEC). As some parameters were missing for 11 populations (supplementary data S1), we carried out a PCA of all complete cases (65 populations) whose first two axes cumulatively explained 58% of the variation in the elemental composition of soil. We further used the variables for which we had complete data (pH H_2_O, pH KCl and bioavailable Ca) as explanatory variables to impute the missing values of the remaining elements by fitting a linear model with the function *predict* from the R package *stats* v.4.1.2 (R Core Team 2021). We then recalculated the PCA, this time including scaled data for pH, total nutrients content and CEC (Fig. 1c), using the *prcomp* function in the R package *stats* v.4.1.2 (R Core Team 2021) and extracted the scores on the PC1 axis, which we used as the primary soil predictor in all the analyses below.

### Genome-resequencing, variant calling, filtration and repolarization

The 76 sequenced populations consisted of 1 to 13 individuals, 6 on average, with an average depth of coverage of 25×. Reads of low quality and adaptors were trimmed and filtered using trimmomatic-0.36 (Bolger et al. 2014) and refined reads were then mapped to a reference genome of North American *A. lyrata* (Hu et al. 2011) using *bwa* v.0.7.15 (Li and Durbin 2009) with the default setting. Duplicate reads were marked using *picard* v.2.8.1 (https://github.com/broadinstitute/picard) and discarded. Afterwards, we called and filtered genotypes with *Genome Analysis Toolkit* (GATK) v.3.7 (McKenna et al. 2010) according to GATK Best Practices recommendations (DePristo et al. 2011; Van der Auwera et al. 2013) and filtering strategies established in our previous studies involving autopolyploid *A. arenosa* populations (Monnahan et al. 2019; Bohutínská et al. 2021) (the complete variant calling pipeline is available at https://github.com/vlkofly/Fastq-to-vcf). Specifically, we used the *HaplotypeCaller* module, which enables the calling of variants per individual with respect to *a priori* known ploidy (set by the – *ploidy* argument) and the calling of variants aggregated across all individuals by the *GenotypeGVCFs* module, allowing for analyses of mixed-ploidy datasets (DePristo et al. 2011; Van der Auwera et al. 2013). We kept biallelic SNPs that passed the GATK-recommended ‘best practices’ filtering parameters (FS>60.0 || SOR>3 || MQ<40 || MQRankSum<-12.5 || QD<2.0 || ReadPosRankSum<-8.0). Further, we masked genes displaying excessive heterozygosity of at least five fixed heterozygous SNPs in at least two populations as being potentially paralogous, following Monnahan et al. (2019). In addition, we masked sites with excessive depth of coverage defined as sites with depth that exceeded twice the standard deviation of depth of coverage in at least 20 individuals. Further filtering based on depth and missing data was done specifically for particular analyses (detailed below).

To infer the ancestral state of each SNP in the reference genome, we constructed the polarization key using the program *est-sfs* (Keightley and Jackson 2018) with two outgroup genomes and allelic frequencies of the focal species. For the outgroups we used *Arabidopsis thaliana* and *Capsella rubella*, and we merged the homologous sites of the outgroups and the *A. lyrata* reference by genomic alignment of all species. Allelic frequencies of the focal species (*A. arenosa*) were parsed by randomly sampling 56 chromosomes from the complete dataset. *Est-sfs* infers the probability of an allele being ancestral, and we categorized alleles as ancestral if the probability was 0.7 or greater. The scripts that were used to generate the polarization key are available at https://github.com/vlkofly/Arabidopsis-genetic-load/tree/main?tab=readme-ov-file#polarisation-using-multiple-reference-genomes.

### Evolutionary relationships between populations

To reconstruct the population genetic structure and relatedness and to investigate diversity, we applied several tools using putatively neutral fourfold degenerated (4dg) biallelic SNPs with the maximum fraction of missing individual genotypes set to 0.5 and a minimum depth of 8× (1.347.998 SNPs, 25× average depth and 12.19% missing data). As a first step, we ran principal component analysis (PCA) on individual genotypes using the *glPcaFast* function (https://github.com/thibautjombart/adegenet/pull/150) in R, reading in the first 10,000 SNPs (see supplementary fig. S2). We also investigated relationships between populations of each ploidy using an allele frequency covariance graph produced using *TreeMix* v.1.13 (Pickrell and Pritchard 2012) (supplementary fig. S3 and Fig. 1d,_e). Input files were created using Python scripts available at https://github.com/mbohutinska/TreeMix_input, filtering the vcf files to achieve a maximum fraction of missing individual genotypes equal to 0.2. We rooted the trees using as an outgroup a diploid population of *A. arenosa* (BDO, Budaörs, Hungary) from the Pannonian lineage, which is the most ancestral lineage of the species (Kolář et al. 2016). We bootstrapped the trees’ topology using a bootstrap block size of 1 kbp (the same window size as used for the selection scan; see below) and 100 replicates and then visualized the results using *FigTree* v.1.4.4. For the full dataset for each ploidy, we repeated the analysis over the range of 0 to 5 migration edges to investigate the change in the explanatory power of the model when assuming inter-population gene flow (supplementary fig. S3). Further, we calculated within-population metrics (nucleotide diversity (π), Tajima’s D (Tajima 1989), downsampling each population to 4 individuals at each site in order to reduce bias due to different sampling size using custom python scripts implemented in *ScanTools*, available at https://github.com/sonia23cld/ScanTools_ProtEvol – window size: 50,000, minimum SNPs per window: 10. With the same method we calculated Rho—an *F*_ST_ alternative designed to be comparable between ploidies (Ronfort et al. 1998; Meirmans et al. 2018)—for each population pair of our dataset. Finally, we reconstructed the population structure and inferred ancestry parameters using a hierarchical Bayesian method tailored to mixed-ploidy datasets, implemented in *Entropy* v.2.0 (Shastry et al. 2021). As input, we used pruned filtered 4dg vcf files (6,942 SNPs, 0.9% missing data) and, as a first step, we applied *k*-means clustering of the results of linear discriminant analysis of estimated genotype likelihoods, using the *MASS* package v.7.3-57 (Venables WN and Ripley BD 2002) in R. This aims to generate initial admixture proportions which then further aid Markov chain Monte Carlo (MCMC) convergence. We then executed Entropy for *K* ranging from 3 to 6, with 60,000 MCMC steps, 20,000 burn-in steps discarded, thinning interval of 20 steps after burn-in. We visualized the genotype estimates in R (https://github.com/sonia23cld/Entropy).

A complete overview of the genomic data used for these analysis and the filtrations applied can be found in supplementary table S1.

### Detecting the signature of selection and candidate adaptive genes

We designed three complementary approaches which conservatively allowed us to identify candidate genes responsible for adaptation to the gradient of siliceous and calcareous soil conditions. First, we applied a window-based scan for directional selection leveraging a manifold-replicated natural set-up to find genetic regions which show a consistent footprint of divergent selection between siliceous and calcareous populations across the species’ range. Keeping into consideration differentiation in soil parameters (PC1 scores in PCA calculated from soil data), but also genome-wide differentiation (reflected by the pairwise *F*_ST_ index; the *F*_ST_ metric is appropriate for comparisons within ploidies), geographical distance and the number of available sequenced individuals, we designed 7 couples of diploid and 7 couples of tetraploid populations (paired dataset – Fig. 1) to be used as contrasts for divergence scans. Importantly, neutral pairwise genome-wide differentiation between calcareous and siliceous populations (4dg-Rho) of the paired dataset is very close between the ploidies (mean 2x: 0.20, SD: 0.05; mean 4x: 0.13, SD: 0.06), indicating a comparable neutral divergence between siliceous and calcareous populations across ploidies. For estimating local divergence, we focused on *F*_ST_ because it is a standard metric used in divergence scans. In later inter-ploidy comparisons, we used gene lists, not absolute values, thus avoiding the issues arising from comparing *F*_ST_ values across ploidies (Meirmans et al. 2018). We used the ScanTools pipeline and calculated pairwise *F*_ST_ for non-overlapping 1-kbp windows along the genome, excluding potential non-informative windows with less than 10 SNPs, as previously applied in diploid and autotetraploid populations of *A. arenosa* (Bohutínská et al. 2021; Konečná et al. 2021). We then identified the 1% quantile of SNPs with the highest *F*_ST_ values in each population pair and annotated them to genes using *A. lyrata* v.2 gene annotation (Rawat et al. 2015) (genes including 5′ untranslated regions, abbreviated as 5′ UTRs, start codons, exons, introns, stop codons and 3′ UTRs). The final lists of top candidate genes included then only those candidates that were identified as above in at least two population pairs, in each ploidy separately. We used a conservative approach, further considering only those gene candidates that were identified as shared between population pairs representing distinct genetic lineages inferred as supported branches by Treemix (these lineages are assigned a different name in Fig. 1e,_f and supplementary fig. S3) and distinct genetic clusters in Entropy (supplementary fig. S4). We tested the magnitude of gene overlap in the final list as non-random using Fisher’s exact test in the R package *SuperExactTest* v.1.1.0 (Wang et al. 2015). In addition, we tested if these candidate genes fall in regions of particularly high recombination, using genome-wide recombination rate values estimated for *A. arenosa* diploids by (Dukić and Bomblies 2022) and performed a one-sided permutation test with 10,000 permutations in R.

Second, we further analysed the results of the selection scan through the *PicMin* pipeline (Booker et al. 2023), a novel method using order statistics to estimate the significance of parallel molecular evolution over three or more lineages (available at https://github.com/TBooker/PicMin). Here, in order to correct for non-independent evolution of population contrasts from within the same lineage, we selected only one contrast per lineage (that with the lowest between-population *F*_ST_, except for pair 12, which had a soil parameter differentiation value greater than pair 14), resulting in a total of 6 lineages in diploids (all contrasts except pair 7) and 4 in tetraploids (keeping contrasts 8, 9, 11 and 12). We ran *PicMin* with _Adapt set to 0.05 (appropriate for a test of 6–4 lineages) and 10,000 repetitions. We then corrected the output p-values for the false discovery rate (FDR, *q*) and kept only windows with *q* smaller than 0.05 for further gene annotation as above.

Finally, we associated genomic variation of the 74 populations for which soil parameters were available by performing environmental association analysis using LFMM2 (Caye et al. 2019) for each ploidy separately (33 diploid and 41 tetraploid populations). We first filtered the vcf file for depth greater than 8, MAF > 0.05, 10% missing data and retained 1,298,394 SNPs. To single out the best value of *k*, which the model uses as latent factor, we performed a principal component analysis of the genotypic data in R, using the function *prcomp* from *stats* package v.4.2.2 (R Core Team 2021), followed by *k*-means clustering (supplementary fig. S5). Based on these results, we applied *k*=4 for each ploidy dataset. We tested the association of the average allele frequencies at each SNP (1.98% missing data were imputed) for each population with the original soil conditions (approximated by the soil PC1 score). We then identified significant soil-condition-associated SNPs by transforming the resulting calibrated p-values with false discovery rate (<_0.1), using the R package *qvalue* v.2.26.0 (http://github.com/jdstorey/qvalue) and annotated the SNPs into genes as explained above. We retained only those genes for which we identified at least three significant SNPs. The final list of top candidate genes used for the following analysis was obtained by retaining all genes that overlapped between the 1% outliers in the *F*_ST_ selection scan and at least one of the other two methods explained above (PicMin and LFMM – Fig. 2). To infer the functional involvement of our candidate genes in potential biological processes, we worked with *A. thaliana* orthologs of *A. lyrata* genes obtained using *biomaRt* (Durinck et al. 2009) and the *A. thaliana* gene universe to perform a Gene Ontology (GO) enrichment test, using the R package *topGO* v.2.46.0 (Alexa A 2024). We applied the *weight01* algorithm with Fisher’s exact test (p < 0.05) as implemented by the package.

A complete overview of the genomic data used for these analyses and the filtering applied can be found in supplementary table S1.

### Selective sweep magnitude and breadth

Differences in the visual profiles of selective sweeps between ploidies have previously been described in studies based on genomic simulations (Monnahan and Brandvain 2020). Here we tested these assumptions on natural data. As the species Arabidopsis arenosa harbours very high levels of segregating variation (Arnold et al. 2015; Monnahan et al. 2019), a selective sweep is not clearly visible when Tajima’s D metrics or other within-population diversity metrics are used, especially considering the number of samples available. For this reason, we focused on between-population differentiation, calculated as Rho for each population pair designed as above. We calculated allele frequencies and Rho metrics for the paired dataset using the ScanTools pipeline onto a vcf file containing data on all biallelic SNPs (supplementary table S1; the output was used also for the calculations presented in Fig. 3a, b,_and d). For each top candidate gene, we measured its selective sweep in terms of magnitude and breadth. ‘Magnitude’ was calculated as the difference between the highest Rho value within the sweep region and the average genome-wide Rho for that population pair. ‘Breadth’ was calculated by extracting Rho values for 100 kbp at each side from the middle of the gene (i.e. the curve region) and fitting a curve using the function loess.as from the R package fANCOVA v.0.6-1, which automatically sets a smoothing parameter using the bias-corrected Akaike information criterion (AICC). Specifically, ‘breadth’ was calculated as the length of the region, in bp, under the sweep curve, where the borders corresponded to the intersection of the fitted curve with the defined threshold of double the value of ‘background’ Rho (average Rho in the 200 kbp flanking the curve region).

### Identifying the evolutionary source of shared adaptive variants

After having identified a shortlist of parallel candidate genes (top candidate genes) likely associated with adaptation to siliceous or calcareous soil in *A. arenosa*, we proceeded to infer their evolutionary source. Using a designated composite likelihood-based method, referred to as the ‘distinguishing among modes of convergent adaptation’ (DMC) (Lee and Coop 2017; Lee and Coop 2019), we modelled patterns of (i) neutral variation, (ii) parallel selection operating on *de novo* mutations at the same loci, (iii) parallel selection acting on pre-existing standing variation, and (iv) transfer of variants through gene flow. For each scenario, this method compares patterns of allele frequency covariance of deviation from the ancestral state (the coancestry coefficient) at loci near the site subject to selection, within and between (selected and non-selected) populations, as a function of the recombination rate. Because this approach is based on allele frequency covariance, which is a relative measure rather than an absolute one, it remains informative regardless of ploidy level. Whereas autotetraploids exhibit additional complexities due to polysomic masking and potential dosage effects, the core principles of coancestry-based inference remain valid. By running DMC in a loop (https://github.com/mbohutinska/DMCloop) over all top candidate regions, we estimated the composite log-likelihoods of each scenario for each parallel gene along each of the 21 population quartets (15 quartets in diploids and 6 in tetraploids), using the same pairs as for the *PicMin* analysis. We set the model parameters taking into account the populations’ evolutionary history inferred by our previous investigations in *A. arenosa* (Bohutínská et al. 2021; Konečná et al. 2021), indicating no difference between the ploidies; see supplementary table S2. Composite log-likelihoods were calculated at twelve locations at equal distances along each gene (plus 25 kbp upstream and downstream), as recommended by the author at https://github.com/kristinmlee/dmc and in accordance with the range of LD decay (150–800 bp) previously estimated in *A. arenosa* (Bohutínská et al. 2021). Not knowing the exact direction of the habitat shift for each quartet, we calculated results for the divergence caused by selection in populations growing on both siliceous and calcareous soil and merged the results. We then identified the best-fitting scenario for each gene and each quartet, only accepting significantly non-neutral cases inferred by comparing models with simulated neutral data in a similar manner as in Bohutínská et al. (2021).

### Construction of protein–protein interaction networks

Functional protein-protein interaction networks (PPIN) were constructed using nodes and edges downloaded from the STRING v.12 (https://string-db.org) (Szklarczyk et al. 2023) protein network database (scored links between proteins) for *A. thaliana* proteins. We did not subset node interactions by applying a threshold on the protein–protein interaction combined score; however, our results do not change when it is applied (supplementary table S3). We associated the ID of each protein sequence (Arabidopsis Genome Initiative, AGI) to a UniProt accession, using the TAIR10 release of Ids conversions (https://www.arabidopsis.org/download/list?dir=Proteins%2FId_conversions) (Berardini et al. 2015). Within the lists of top candidate genes, two and three genes were not included in the construction of the PPIN in diploids and tetraploids, respectively, as they could not be associated to any *A. thaliana* ortholog. Moreover, for one gene in each ploidy there was no associated protein in the STRING database, leaving us with a final number of 40 and 24 candidate genes in diploids and tetraploids, respectively. For each ploidy dataset, we tested if the average node number of interactions (connectivity) was significantly greater than the average node connectivity of the full *A. thaliana* dataset by performing a one-sided permutation test with 10,000 permutations in R. We then visualized each ploidy PPIN in R using the packages *igraph* v.1.3.1 (Csardi and Nepusz 2006) and *tcltk* v.4.1.2 (Grosjean 2022).

## Acknowledgements

We thank Miroslav Poláček and Karolína Havlíková for assistance during fieldwork and Magdalena Bohutínská, Levi Yant, Edita Tylová, Stanislav Kopřiva, Patrick Monnahan and members of the Plant Ecological Genomics group for feedback on an earlier version of this manuscript.

This research was funded by the Charles University (GAUK project 243-252135 awarded to SC), the European Research Council (ERC) under the European Union’s Horizon 2020 research and innovation programme (ERC-StG 850852 DOUBLE ADAPT to FK), and the Czech Science Foundation (project 23-07204M to FK). Additional support was provided by the Czech Academy of Sciences (long-term research development project RVO 67985939). Access to computing and storage facilities owned by the parties and projects contributing to the National Grid Infrastructure MetaCentrum was provided under the programme ‘Projects of Large Research, Development, and Innovations Infrastructures’ (CESNET LM2015042) and is greatly appreciated.

## Notes

### Competing Interest Statement

The authors have declared no competing interest.

