## supplementary fig. for "Whole-genome duplication reshapes adaptation: autotetraploid *Arabidopsis arenosa* leverages its high genetic variation to compensate for selection constraints"

**Supplementary figures and tables:**

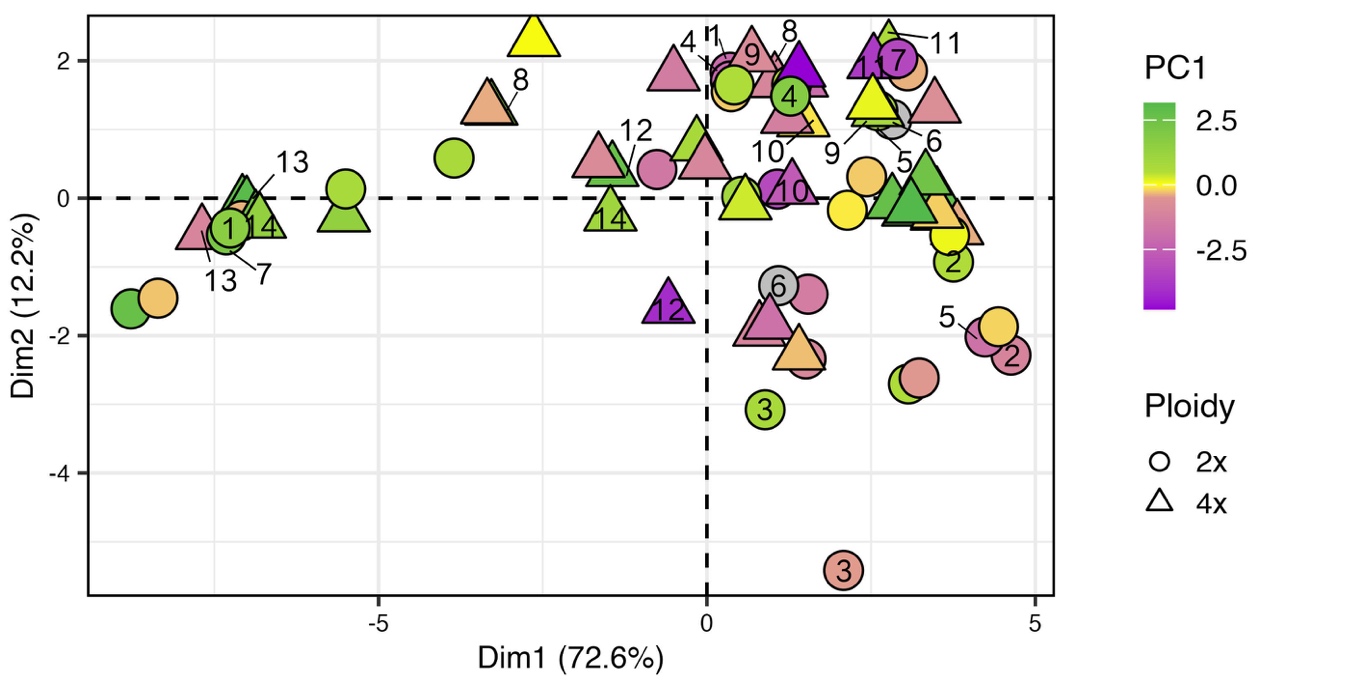

**Figure S1.** PCA of the 19 bioclimatic variables extracted for locations of each population of the full dataset from WorldClim database (<https://www.worldclim.org/data/bioclim.html>). Populations used in the paired dataset are indicated by their corresponding number. Green-violet color ramp reflects the position along the PC1 axis of PCA analysis of soil parameters at the population location of origin. Shape refers to the ploidy of the populations. Note the lack of obvious association between environmental conditions and either soil conditions or ploidy.

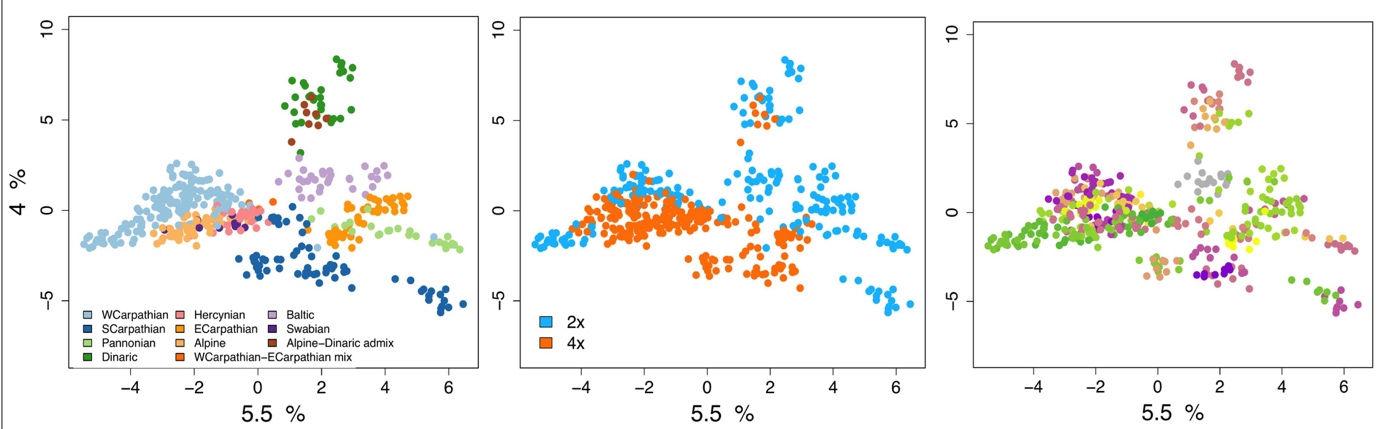

**Figure S2.** PCA of individual genotypes calculated from 10,000 neutral fourfold degenerated (4dg) biallelic SNPs. From left to right individuals were colored by lineage, ploidy and position along the PC1 axis of soil parameters at the population location of origin.

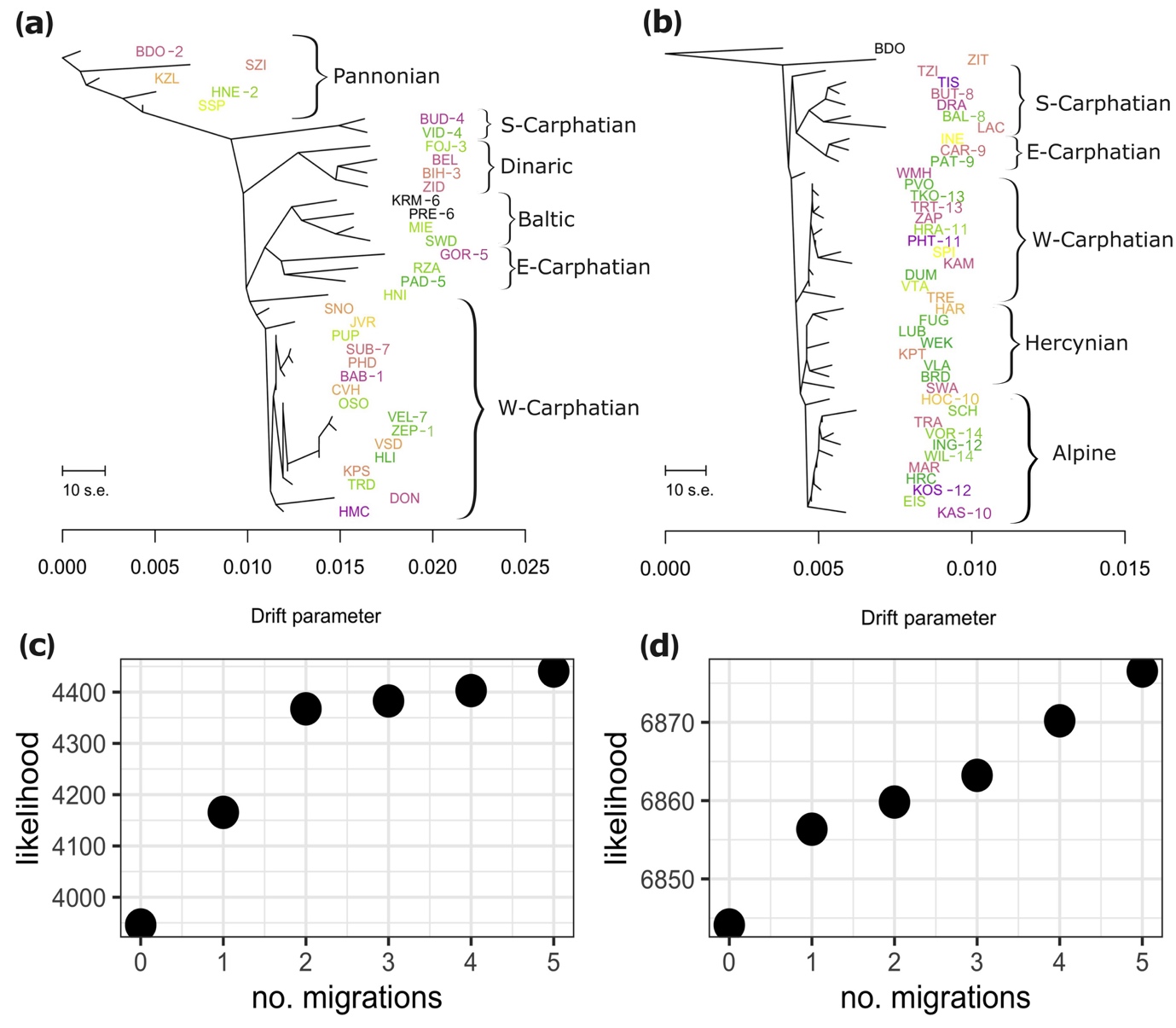

**Figure S3.** a, b) Relationships between the diploid (a) and tetraploid (b) populations of the full dataset depicted using an allele frequency covariance graph calculated in Treemix (1,347,998 biallelic SNPs). Populations used also in the paired dataset are labeled by their corresponding pair number. Groups of branches are annotated by major genetic lineages. Green-violet color ramp reflects the position along the PC1 axis of PCA analysis of soil parameters at the population location of origin ranging from calcareous (violet) to siliceous (green).

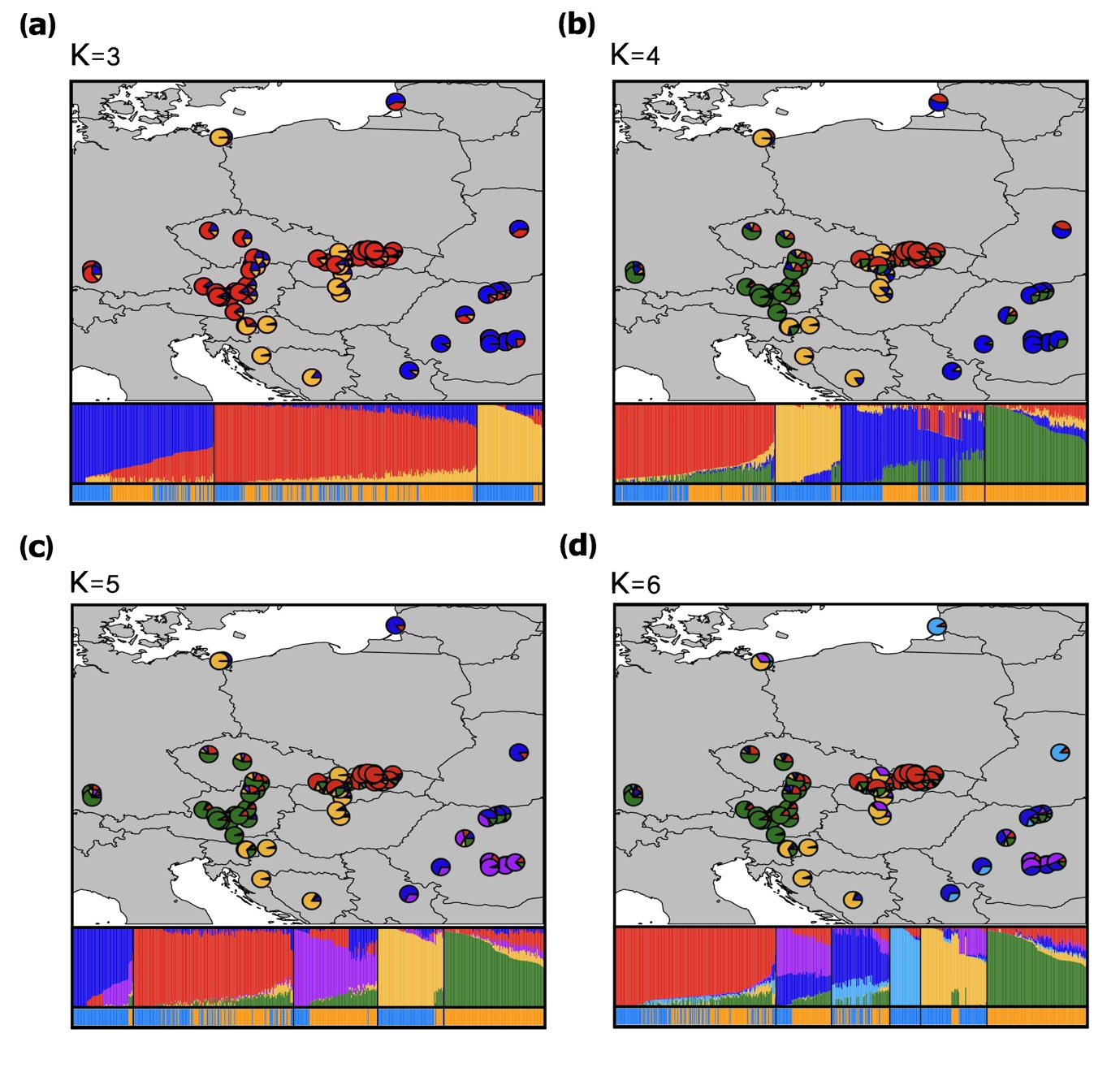

**Figure S4.** Range-wide genetic differentiation of *A. arenosa* full dataset inferred by means of population clustering. In each picture we show the geographical distribution of the populations indicated as pie charts reflecting the proportional assignment to specific genetic clusters identified by Entropy, individual assignment into each genetic cluster identified by Entropy and individual’s ploidy (lightblue – diploid, orange – tetraploid). Here we show the results for a defined number of clusters (K) of three (a), four (b), five (c) and six (d). In line with TreeMix analysis, Entropy results for k=6 identify five and four main clusters/lineages for diploid and tetraploid respectively: Alpine/Hercynian – 4x (green), Western-Carpathians 2x-4x (red), Dinaric/Pannonian 2x (yellow), Southern-Carpathians 2x-4x (purple), Baltic 2x (lightblue), Eastern-Carpathian 2x-4x (blue). Note that although tetraploids had originated singularly from the Western Carpathian diploid lineage (Arnold et al. 2015; Monnahan et al. 2019), local populations from the three mixed-ploidy regions (Western-Carpathians, Southern-Carpathians, Dinaric/Alpine tend to cluster close with locally co-occurring diploids, in line with previous reports of pervasive local interploidy gene-flow (Monnahan et al. 2019).

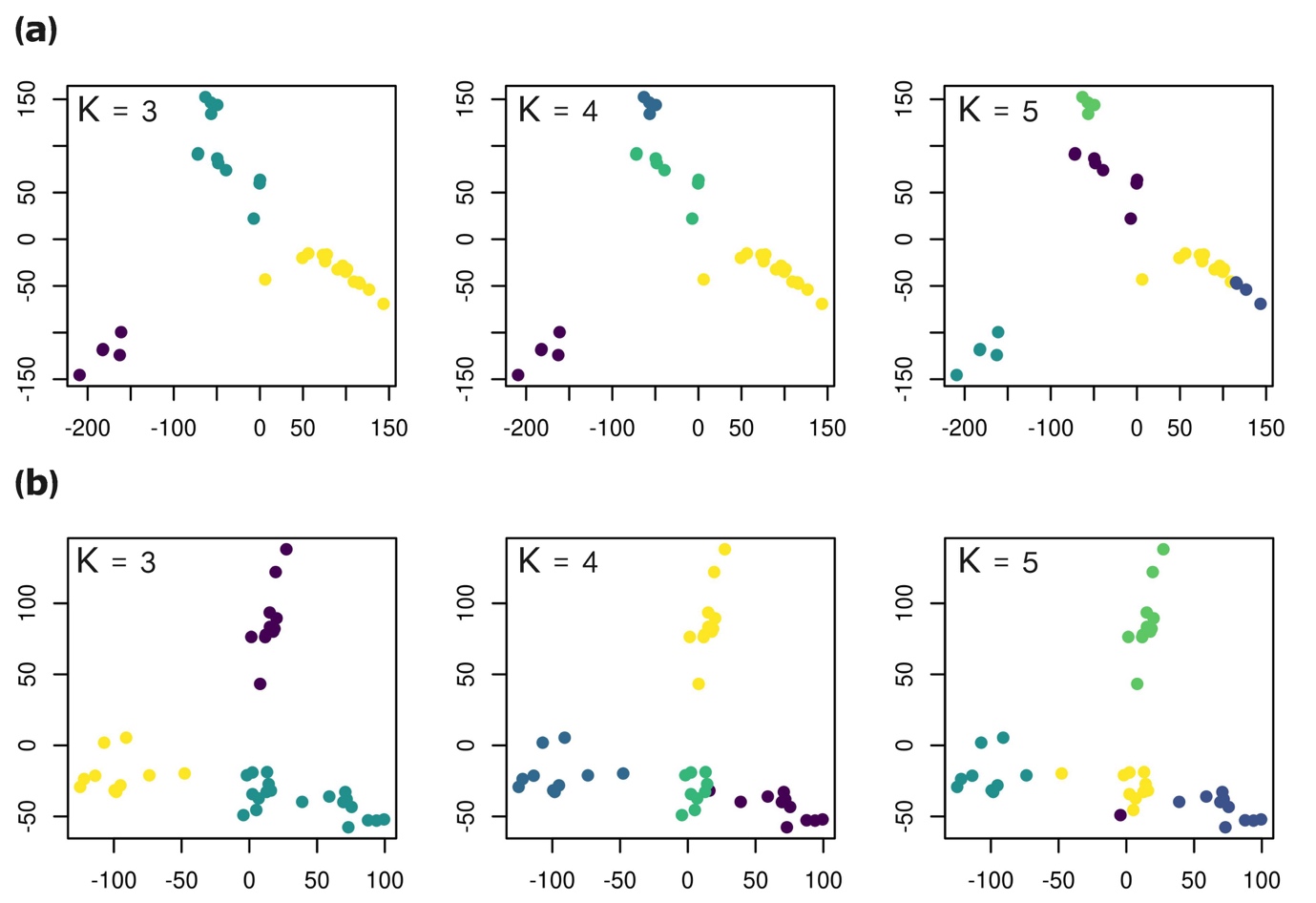

**Figure S5.** a, b) K-means clustering of genetic data of diploid (a) and tetraploid (b) populations (all SNPs) used to perform environmental association analysis using LFMM2. The analyses leaded us to choose K=4 as latent factor in LFMM for both ploidies, which is in line with Entropy results for K=6 (four clusters per ploidy as Baltic 2x lineage is not included in the LFMM dataset).

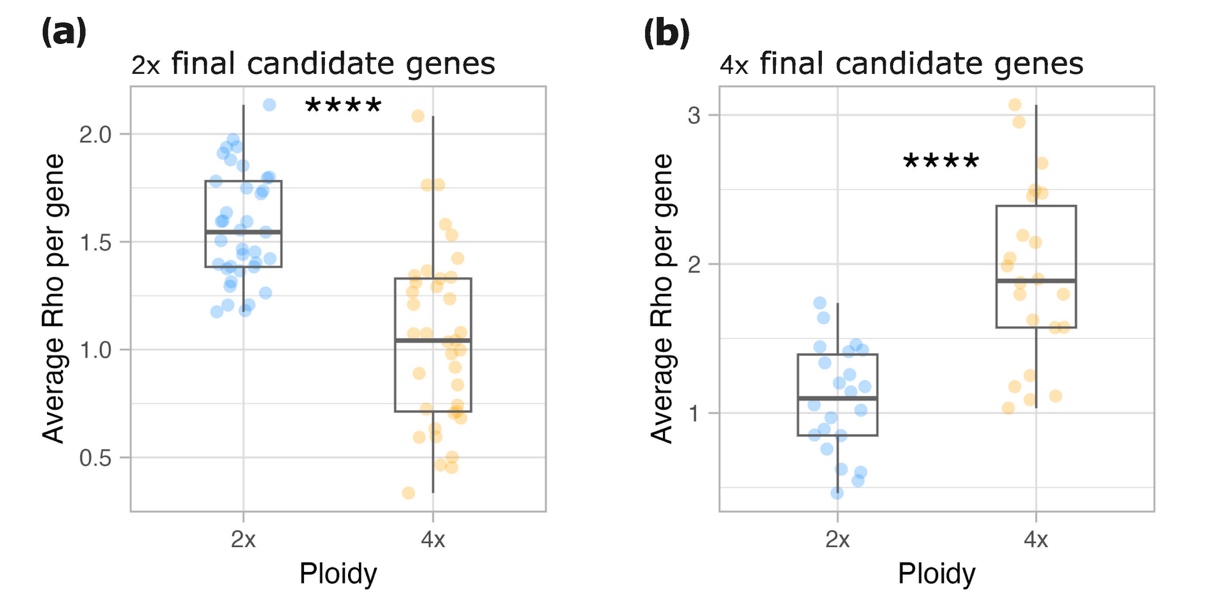

**Figure S6.** Average increase in genetic differentiation (Rho) relatively to genome-wide values reciprocally calculated in both ploidies for diploid final candidate genes (a) and tetraploid final candidate genes (b) across all populations pair of the paired dataset (one dot is one gene average Rho across all population pairs). The six final candidate genes shared between ploidies were removed from the calculations to avoid pseudo-replication. In both scenarios, Rho is significantly higher in the ploidy dataset where the candidate genes are under positive selection (candidates) – scenario a: t(60.331) = 6.4695, p-value = 1.953e-08; scenario b: t(34.512) = -5.6264, p-value = 2.51e-06, Welch Two Sample t-test.

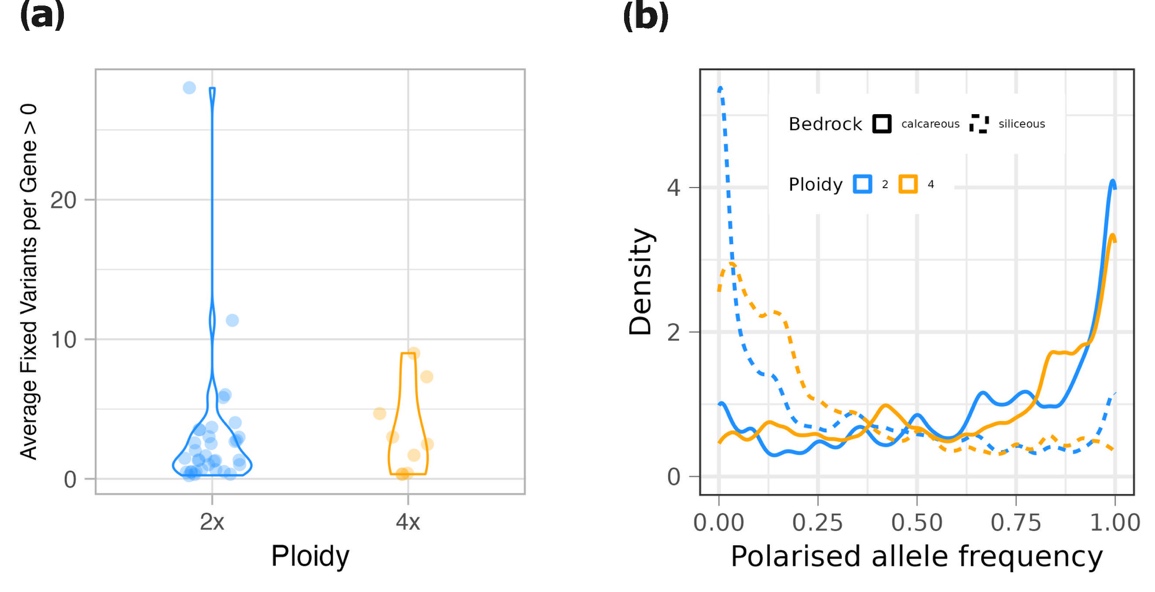

**Figure S7.** Different fixation rates in top candidate genes for substrate adaptation in diploid and tetraploid *A. arenosa* populations when sampling an equal number of chromosomes (12) in each population (paired dataset) a) Average number of fixed variants per top candidate gene calculated across overlapping 1-kbp sliding window and population pairs. While 34 top genes in diploids show at least one fixed SNP (79.07% of the total), in tetraploids only 9 (32.14% of the total) top genes show fixation across all population pairs. b) Distribution of the average allele frequency per SNP (polarised by soil type to aid visualisation in the functional context) across ploidies and soil types is more shifted towards intermediate values in tetraploids. Average allele frequency was calculated per each SNP belonging to a top candidate gene, with flanking region of 2-kbp. In both a and b, each pair was added to the calculation of the mean only if the gene was a significant outlier in that specific pair.

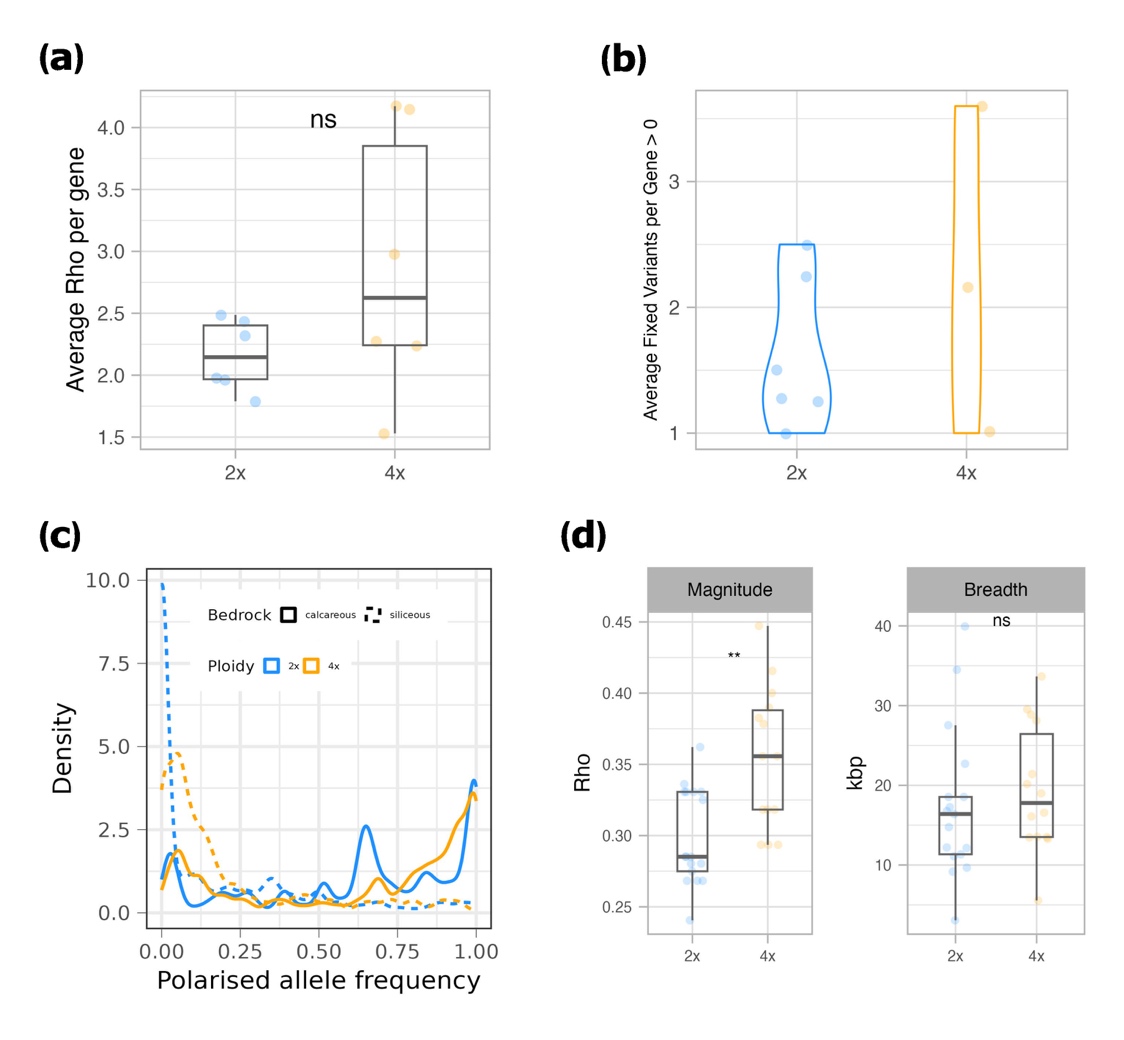

**Figure S8.** Different footprints of selection for substrate adaptation in the six candidate genes shared between ploidies (paired dataset). a) Average increase in genetic differentiation (Rho) for each shared gene relatively to genome-wide values. b) Average number of fixed variants for shared gene calculated across overlapping 1-kbp sliding window and population pairs. In both a and b, each pair was added to the calculation of the mean only if the gene was a significant outlier in that specific pair. c) Distribution of the average allele frequency per SNP (polarized by soil type to aid visualization in the functional context) across ploidies and soil types. Average allele frequency was calculated per each SNP belonging to a shared candidate gene, with flanking region of 2-kbp. Only candidates from their relevant population pairs where they appeared as outliers were included in the calculations. d) Sweep magnitude and breadth for each shared candidate gene across relevant population pairs.

**Table S1.** Summary of datasets and filtrations for genomic analyses.

| **Analysis** | **Dataset** | **Filtrations** | **N. Populations** |
| --- | --- | --- | --- |
| **PCA** | 4dg biallelic SNPs | 10.000 SNPs | 76 |
| **TreeMix, Pairwise Fst and Rho, nucleotide diversity, Tajima’s D** | 4dg biallelic SNPs | MFFG 0.2 and DP < 8 at the population level | 76 |
| **Entropy** | 4dg biallelic SNPs | MFFG 0.2 and DP < 8, pruning over 1kbp windows and 100 bp distance between windows, minf 0.05, maxf 0.95 | 76 |
| **Fst selection scans,**  **Rho candidate genes** | all biallelic SNPs | MFFG 0.2 and DP < 8 at the population level | 14 (2x) – 14 (4x) |
| **PicMin** | all biallelic SNPs | MFFG 0.2 and DP < 8 at the population level | 12 (2x) – 8 (4x) |
| **LFMM2** | all biallelic SNPs | MFFG 0.1, DP < 8,  MAF 0.1, imputation missing data | 74 |
| **Gene Heatmap and Local PCA** | all biallelic SNPs | MFFG 0.1, DP < 8,  MAF 0.1, imputation missing data | 76 |
| **DMC** | all biallelic SNPs from selected genes | MFFG 0.2 and DP < 8 | 12 (2x) – 8 (4x) |
| **DMC (neutral data)** | 4dg biallelic SNPs | MFFG 0.2 and DP < 8 | 12 (2x) – 8 (4x) |

4dg sites: fourfold-degenerate sites

DP: read depth

MFFG: max fraction of missing individual genotypes

minf: minimum allele frequency

maxf: maximum allele frequency

MAF: minor allele frequency

Before specific filtrations, 4dg vcf file of full dataset was filtered for MFFG 0.5 and DP < 8 and contained 1.347.998 SNPs, 12.19% missing data.

**Table S2.** Parameters used to perform convergent adaptation models in DMC.

| **Parameter** | **Parallel selection models** | **Ploidy** | **Values used** |
| --- | --- | --- | --- |
| Selection coefficient | All models | 2x, 4x | 0.0001, 0.001, 0.01, 0.05, 0.1, 0.2, 0.5 |
| Migration rate | Migration | 2x, 4x | 1e-07, 1e-06, 0.00001, 0.0001, 0.001, 0.01 |
| Generations since populations split | Standing variation | 2x, 4x | 100, 1000, 5000, 10000 |
| Initial allele frequencies prior to selection | Standing variation | 2x, 4x | 2.5e-06, 0.00001, 0.0001, 0.001, 0.01 |

Effective population size Ne = 100,000

Per base pair recombination rate of 3.7e^-8^

**Table S3.** Result parameters for each protein-protein interaction network (PPIN) built for each ploidy dataset applying several levels of combined score thresholding.(Booker *et al.* 2023)

| **Score thr.** | **Ploidy** | **N. genes** | **Mean** | **Min** | **Max** | **P-value** |
| --- | --- | --- | --- | --- | --- | --- |
| 300 | 2x | 39 | 140.2 | 5 | 862 | 0.6192 |
| 300 | 4x | 24 | 268.33 | 13 | 1687 | 0.0202 |
| 400 | 2x | 39 | 80.85 | 1 | 563 | 0.6888 |
| 400 | 4x | 24 | 170.12 | 10 | 1178 | 0.0218 |
| 500 | 2x | 37 | 46.22 | 2 | 210 | 0.8583 |
| 500 | 4x | 24 | 122.3 | 5 | 983 | 0.0282 |

Score thr: protein-protein interaction combined score used as a threshold.

N.genes: number of node genes retained after threshold subsetting.

Mean, Min, Max: mean, minimum and maximum number of interactions.

P-value: one-sided permutation test p-value.
